# Impact of Driving Dynamic Fluctuations of 6DOF-SVC Model Predictions Across Five Real-World Driving Studies

**DOI:** 10.64898/2026.09.18.752352

**Authors:** Benedikt Buchheit, Yannick Robin, Elena N. Schneider, Mohamad Alayan, Daniel J. Strauss

## Abstract

Despite the use of technological countermeasures for motion sickness (MS), studies show that the sensory conflict, and thus symptoms of MS such as nausea, cannot be completely prevented while traveling as a car passenger. Accordingly, reliable prediction of MS will represent a possible solution for intelligent vehicles to avoid the abortion of the drive, e.g., through adapted driving strategies or route planning. Although the modeling of individual subjective well-being is becoming increasingly important, the prediction of overall motion sickness incidence (MSI) in groups along a route remains a complementary assessment metric. In the present study, application-oriented parameter sets for the popular 6DOF-SVC model to estimate MSI were validated using five previously conducted real-world driving experiments. The results demonstrate that the MSI of unknown routes can be successfully predicted in an application-oriented manner; however, inaccuracies in the predicted MSI trajectories remain. In addition, for the first time, it is shown that the natural MSI fluctuations, based on the driving dynamic profile, route characteristics, or driving style, is approximately 10%. Moreover, findings indicate that individual MSI thresholds lie in a range of 16.65% to 36.73% and slightly correlate with MS susceptibility. However, their high variability across driving scenarios limits their applicability.

## 1 Introduction

When drivers hand over control to a self-driving car, they become passengers, which increases the likelihood of experiencing Motion Sickness (MS) [1], [2]. Symptoms, such as nausea or headaches can arise [3], making it difficult to engage in Non-Driving-Related Tasks (NDRTs), like reading a book. This reduces the benefit of using self-driving cars for leisure activities, work, or entertainment and also reduces users’ acceptance of such driving functions. Avoiding excessive vehicle dynamics, e.g., by operating the vehicle at low speeds, is socially as unacceptable as the daily intake of anti-nausea drugs, which may also cause side effects such as drowsiness or blurred vision [4]. Consequently, there is strong interest in mitigating MS through technical solutions, such as on-board Countermeasures (CMs), that have the potential to ensure user acceptance of automated driving functions.

The most likely primary trigger of MS is a Sensory Conflict (SC), which refers to a discrepancy between expected and actual sensory inputs, particularly those associated with the vestibular system [3], [5], [6]. Therefore, a common technological CM concept is to influence human sensory processing and expectation. Although using anticipatory cues, e.g., indication of upcoming driving maneuvers, or sensory stimulation or substitution [7], [8], show a reduction and delay of MS symptoms regardless of the sensory modality (visual [9], [10], auditory [2], [10], [11], or tactile [10], [11], [12], [13]), these effects have so far not been strong enough to prevent MS entirely. The overall efficacy of these interventions remains uncertain. A major concern is that most studies, such as [2], [14], [15], [16], [17] employ within-subject designs, where participants are aware that a CM is being tested on them. This awareness can induce placebo effects, which are known to alleviate MS symptoms [18], [19]. In addition, other physiological influencing factors, such as mental distraction [7], or the MS questionnaire itself [20], can influence MS experience. Moreover, “feel-good” CM, such as relaxing music, perfume, and ventilation applied after the stimulation phase [14], suggest that participants’ psychological disposition can significantly impact the outcomes. Studies employing between-subject designs [10], [21], however, report minimal or no reduction in MS symptoms, regardless of whether visual or auditory stimulation or feel-good CM are used. Another critical factor is the role of training and habituation. For instance, it remains unclear whether visuospatial ability contributes to MS reduction, see findings of [22] and [23].

In addition to the internal sensory processing and conflict mechanisms of the passenger’s sensory perception, the physical effects of the accelerations and rotations occurring during driving can also be influenced. In principle, an extremely conservative or passive driving style could substantially reduce driving dynamics. However, this approach is inherently limited, as individual target parameters, such as arrival time and energy consumption, as well as the overall traffic flow, must be taken into account during driving. Since driving dynamics cannot be entirely avoided, the focus shifts toward optimizing the directions and distribution of the acting forces. It is well established that the spatial orientation of the passenger’s head and body has a significant influence on the occurrence of MS [24], see also [25]. A pilot study employing an actively swinging-seat-system [26] supports this finding. However, practical applications, such as active-tilting vehicles [19] and experience gained from tilting trains [27], have shown that this physical CM does not always prove effective. It also indicates a long-term habituation to conventional vehicle motion patterns, which makes it difficult for passengers to quickly adapt to unfamiliar situations in the vehicle, such as an active suspension systems.

Given that CMs cannot entirely prevent MS regarding the current state of the art, continued modeling and investigation of MS remain essential. Both for estimating the passenger’s current state and for supporting avoidance strategies, such as route selection or maneuver planning. One of the most established, data-driven MS models for predicting (inner-vestibular) SC is the Six Degress-of-Freedom Subjective Vertical Conflict (6DOF-SVC) model [28], [29]. This model uses the mathematical modeling of the semicircular canals (SCC) and otolith organs (OTO) [30], which compares the three-dimensional movement experienced by the head with an inner ear reference model to estimate the SC [28], [31] based on the subjective vertical conflict (SVC) theory. Based on the fundamental research findings of [32], [33] on the relationship between acceleration attitude and low-frequency vertical oscillation, the 6DOF-SVC model predicts the Motion Sickness Incidence (MSI) of a group of subjects. The 6DOF-SVC model is able to predict MSI ratios of different head and torso movement techniques, which influence MS occurrence [26], [28]. Additionally, it can reproduce experimental results from real-world driving experiments [28] as well as studies with an adapted parameter set simulating a passenger engaged in a NDRT [29]. Since then, the 6DOF-SVC model has been extended with sub-modules, e.g., transfer the vehicle dynamics to the passengers’ head as model input [29] or to estimate the vestibulo-ocular reflex as output [34]. There have also been attempts to extend a version of the 6DOF-SVC model with a *learning and prediction* sub-model to simulate passengers’ internal expectation [35] or to incorporate compensation information from the visual vertical prediction [36], [37], [38], see also [39], and the optimization and modeling of visual MS mitigation [40] to capture variations in vision conditions. However, MSI does not allow for conclusions at the level of individual passengers, as it is derived from group-level data. For this reason, the MSI-computing component of the 6DOF-SVC model was replaced with a nausea sub-model [41], [42] in order to predict the subjective MS experienced by an individual [43]. Since vehicle studies commonly employ the established Misery Scale of MS (MISC), an 0–10 scale for subjectively assessed MS symptoms by the participant [44], [45], to quantify MS, this revised model formulation enables a direct comparison between model predictions and the responses of individual passengers. Although initial approaches appear promising in suggesting that the individualized 6DOF-SVC model can be trained for each passenger [37], [46], [47], its validation across a larger number of driving experiments and diverse driving paradigms is still outstanding.

Individualized modeling of MS is likely to dominate future research and applications. Nevertheless, several limitations and open research questions remain unresolved, which continues to affect the interpretation and reliability of the predictions of the models. One limitation is the subjective nature of symptom manifestation and assessment itself. Although the MISC has proven useful in real-world driving studies, see [7], [12], [15], [45], [48], many experimental paradigms yield low (*≤* 4) average MISC scores, see [2], [7], [13], [15], [49]. It is questionable whether participants can reliably assign their symptoms to specific scale values, e.g., the difference between MISC of 2 (vague dizziness) to 3 (some dizziness), in a single session, especially when placebo effects may unintentionally influence their responses.

Consequently, the signal-to-noise ratio within the subjective rating or MISC signal remains undefined. Potential discrepancies between subjective well-being and reported symptoms have already been discussed [50]. Although findings [45] indicate that the MISC scale is suitable for assessing MS in vehicle-based studies, the variability of form of the day of participants and the reproducibility of driving experiments have not yet been clearly defined. To date, no study has systematically investigated multiple individuals across temporally distinct experimental sessions. Beside that, computing and analyzing the grand average from an ordinal scale, such as the average MISC, poses a methodological challenge. It is not discussed in the literature that average MISC values may not follow a normal distribution, as subjects typically start at a MISC rating of zero, which can result in a distribution skewed toward zero over time. Accordingly, results [43] based on average MISC trajectories should be re-examined using appropriate statistical methods. Furthermore, the temporal dependence of individual query points in the MISC trajectory requires adjustments to the significance level [51], such as applying false discovery rate correction [52]. Moreover, the question arises as to which methodological approach should be applied to participants who abort the measurement prematurely, and whether the MISC value obtained up to that point should be excluded from the average calculation or kept constant from the moment of discontinuation, see also [11], [15]. As individual measurements, rather than overall study outcomes, have increasingly become the primary focus in MS research, the literature shows a continued refinement and optimization of MS models. However, this development often proceeds without adequately accounting for natural variability, error tolerances, or plausible target ranges, both in model predictions and in the corresponding reference measures. Model optimization is frequently driven toward a single target value, without sufficiently questioning the extent to which this target value inherently fluctuates due to inter- and intra-individual variability.

Although these open questions have not yet been fully resolved, there is little doubt that individualized subjective MS assessments already serves as, and will continue to serve as, an accepted standard for evaluation. Nevertheless, individualized MS prediction cannot be applied in every context, as its applicability is inherently limited in scenarios with frequently changing passengers, such as robotaxis or public transportation, where participant-specific calibration is not feasible. Although the prediction of subjective MS measures (e.g., MISC) is a meaningful and appropriate strategy for route planning [53] and may directly improve passenger comfort, the simultaneous prediction of a generalized MSI reference value, for example by using the 6DOF-SVC model, may provide complementary information. In particular, MSI predictions can be used to assess the baseline MS risk of multiple candidate routes independent of a specific passenger and to identify MS critical route segments. Furthermore, generalized MSI prediction represents a practical alternative for vehicles with limited sensing capabilities that cannot directly monitor their occupants, as the required vehicle motion signals are inherently available. Unfortunately, there are currently no public wide-ranging databases that provide MSI values along with corresponding individual MS indices for various driving routes and vehicle dynamics, which limits the development of MS models and avoidance strategies, too. Furthermore, the 6DOF-SVC model, or MSI prediction, lacks fundamental research in the following aspects, which will be addressed and investigated in this work. In doing so, data from five existing real-world driving experiments is used:

1. The MSI trajectory, as well as the applicability of the model to previously unseen driving routes, remains difficult to validate reliably due to the limited amount of research available in this area. To date, the prediction uncertainty associated with MSI trajectories has not been systematically investigated, limiting the ability to contextualize potential prediction errors. Therefore, in a first step, we optimize our previously published parameter sets [29], which predict correct final MSI values as well as MSI trajectories, using two real-world driving experiments for calibration and three additional independent driving experiments for validation. This approach allows the transferability and robustness of the parameterization to be assessed under previously unseen driving conditions.
2. While the first research question focuses on prediction accuracy, that is, the discrepancy between the predicted MSI for a driving experiment and the corresponding experimental outcome, the second research question addresses an aspect that has received little attention so far: the variability of MSI predictions resulting from natural fluctuations in driving dynamics (used as input of our model). The 6DOF-SVC model is a deterministic model that produces exactly one MSI (MSI trajectory) prediction for a given input signal. Consequently, the predicted MSI always corresponds to a specific driving dynamic profile. In practice, however, a route cannot be represented by a single driving dynamic profile. Repeated traversals of the same route, differences in driving style, traffic conditions, or environmental influences inevitably lead to multiple realizations of the vehicle motion. As a result, a single route may generate a range of model inputs rather than one unique input signal. This implies that the MSI associated with a route may not necessarily be described by a single value, but rather by a range of plausible MSI outcomes. Such a range can be interpreted as an MSI prediction corridor bounded by the minimum and maximum MSI values resulting from naturally occurring fluctuations in the driving dynamics or the given driving styles of the driver. To date, no established database exists that quantifies this variability or provides reference values regarding the expected prediction spread of the 6DOF-SVC model. Therefore, in a second step, the present work investigates how MSI predictions vary when natural fluctuations occur in the driving dynamic profiles (fluctuations between driving dynamic profiles rather than fluctuations within an individual profile) of the five driving experiments. The objective is to quantify the magnitude of these prediction fluctuations and thereby assess whether an MSI-corridor-based interpretation should be considered necessary or beneficial when applying this modeling approach in practice. Ultimately, this analysis aims to determine to what extent MSI predictions should be interpreted as a single deterministic value or as a range of expected outcomes associated with a given route.
3. Although the direct prediction of subjective MS represents a valid and promising approach, individual susceptibility may also be reflected indirectly through MSI-based metrics. The potential for mapping MSI to an individual MS index has not yet been investigated; however, it is conceivable that each individual reaches a characteristic MSI threshold prior to the onset of sickness. Therefore, the present dataset is used to investigate whether such individual MSI thresholds exist at the group level, whether they are associated with the participants’ general MS susceptibility, and whether these thresholds remain stable for participants who took part in multiple driving experiments.

## 2 Materials & Methods

### 2.1 Experimental Driving Studies

Our study comprises five real-world driving experiments (S1-S5) conducted over recent years in real vehicles, each addressing different research questions in the field of MS. Despite their varying objectives, these studies share sufficient methodological and contextual similarities to allow for their integration and pre-processing for use in MS modeling. All experiments were conducted at the same research institution and involved participants from a local population (Central European, university environment). The MS susceptibility across the five participant groups was assessed using the average of individual, pre-experiment questionnaire scores (Q-Score). Here, the Q-Score [54] is derived from responses to eight essential, MS related questions (see Appendix Table 9) and cannot exceed a maximum value of 12, indicating highest MS susceptibility. To minimize potential placebo effects and negative expectancy biases toward the experiment, which have previously been shown to influence MS outcomes [20], see also findings on placebo and expectation effects in MS countermeasures [14], [15], the questionnaire was intentionally kept concise and non-intrusive, as well as, for this reason, no MSSQ was used. While the experiments, the participants were seated in the back seat and had to solve NDRTs continuously under different conditions. The subjective MS level of the participants was collected at regular intervals on a visual version of MISC [44] (0 - no symptoms at all to, 10 - vomiting or abortion of the measurement), which enables a faster and more intuitive rating while driving. When the MISC rating reached 10, the measurement was aborted. Participants who rated MISC of at least 7 are classified as motion sick in our experiments.

In the first experiment (S1) [29], isolated maneuvers (e.g., slalom, stop and go, traffic circle, speed bumps) were carried out in a closed testing field. The overall duration was approximately one hour and included low-dynamic baseline laps, with 18 participants included.

The second experiment (S2) [29], [48] was an one-hour drive through real road traffic, including motorway, interurban, and urban sections, with a total of 20 participants.

In the third experiment (S3) [17], [55], 29 participants completed two trials to test a technological CM in comparison to a non-operational (or placebo) CM, while driving on a closed testing area for 23 minutes. The non-functional CM plays low-frequency random color changes that have no detectable effect on MS occurrence. For the current evaluation, only the non-functional CM trial is included. Compared to the other experiments, the distribution of participants’ MS susceptibility is slightly inhomogeneous, as more susceptible individuals participated, resulting in an increased average Q-Score.

In the fourth experiment (S4), [19], 23 participants were driven twice on the experimental route, once in a standard (reference) vehicle and in a vehicle with active suspension. The route was also on a closed testing area and had a duration of 15 minutes. Similar to the third experiment, only the trial using the reference vehicle is included, because, compared to the active suspension system, it does not affect the MS occurrence. Due to the experimental design and the availability periods of the vehicles, there is a two-week gap between the measurement drives, using the reference vehicle, of the first 11 participants and the remaining 12.

Whereas experiments S3 and S4 were within-subject paradigms, the fifth experiment (S5) [10] employed a between-subjects design, comparing two technological CM groups to a control group (n = 15) without a CM. They drove on a closed testing area for around 20 minutes. Only the control group is included in the data analysis.

In S2, S3, and S5, the test car was a family van (VW Touran, 2^*nd*^ gen.), in S1 a family van (Ford Galaxy, 3^*rd*^ gen.), and in S4 an SUV (Audi Q5, 2^*nd*^ gen.). See experimental designs, driving routes and g-g-diagramms in Appendix Fig. 8, 9, 10.

All experimental procedures complied with the tenets of the Declaration of Helsinki and were approved by the Institutional Review Board at Ethikkommission der Ärztekammer des Saarlandes, Saarbrücken, Germany, Identification Number: 199/17, 181/18, 191/22 and 149/23. Informed consent was obtained from each participant.

Finally, our evaluation focused on five participant groups, Table 1, which developed MS symptoms while driving in a passenger car and performing NDRTs. All 105 participants included in the study were assigned to the control condition, which did not involve any CM or technological intervention. As described above, we have no reason to assume that the selected participants from S3, S4, and S5 were subject to physiological influences that could have impacted the MS occurrence of the driven route. Driven baseline phases, during which participants did not perform the NDRT, were excluded from the recorded driving dynamic profile because participants moved and looked around freely to avoid MS symptoms during these phases. As a result, every analyzed measurement starts at the time the participants/ passengers lowered their gaze to solve the NDRT. No further exclusions were applied to this participant pool, as all datasets required for modeling are fully available on reasonable request. Nine participants took part in more than one study and were analyzed also in an additional analysis step (see Methods).

**Table 1:** Overview about the number of included participants (n), MS occurrence, duration (T) of the ride, and distribution of MS susceptible and resistant participants, for the five driving experiments.

| No. (year) | n | MS occurrence [%] | T [min]* | MS susceptibility distribution (average/ std. Q-Score**) |
|---|---|---|---|---|
| S1 (2017) | 18 | 55.6 | 60.0 | normal distributed (4.88/ 2.89) |
| S2 (2018) | 20 | 45.8 | 51.7 | symmetrically truncated normal distributed (5.13/ 2.92); with-out extreme resistant and susceptible participants |
| S3 (2021) | 29 | 79.3 | 19.4 | truncated distributed (7.31/ 3.20); without resistant partici-pants |
| S4 (2022) | 23 | 45.5 | 14.1 | normal distributed (5.52/ 4.29) |
| S5 (2023) | 15 | 20.0 | 17.8 | symmetrically truncated normal distributed (5.80/ 3.73); with-out extreme resistant and susceptible participants |

\*without baseline
\*\*MS susceptibility index

### 2.2 6DOF-SVC Model

#### Structure of the 6DOF-SVC Model

To predict the MSI for the different experiments the 6DOF-SVC model was applied in the basic form [28], see also [56], see Fig. 1, integrated in MATLAB/Simulink (R2019a, The MathWorks Inc.). Although several extensions of the 6DOF-SVC model have been proposed in recent years, the present analysis is based on the original base model. This decision is motivated by the following considerations: 1) Certain sub-modules, such as those modeling vestibular components [34], [36], [37], [38], cannot be applied to the available database, which comprises sequential, highly mixed, and experiment-specific NDRTs. 2) The baseline model is the most extensively studied and, compared to its extensions, involves a smaller number of parameters, thereby reducing the influence of unknown or insufficiently understood factors. This aligns with the modeling principle that all models should achieve the required level of accuracy with the lowest possible complexity. 3) Our published parameter sets that are validated in this study were originally developed and verified using the base model.

**Figure 1:**
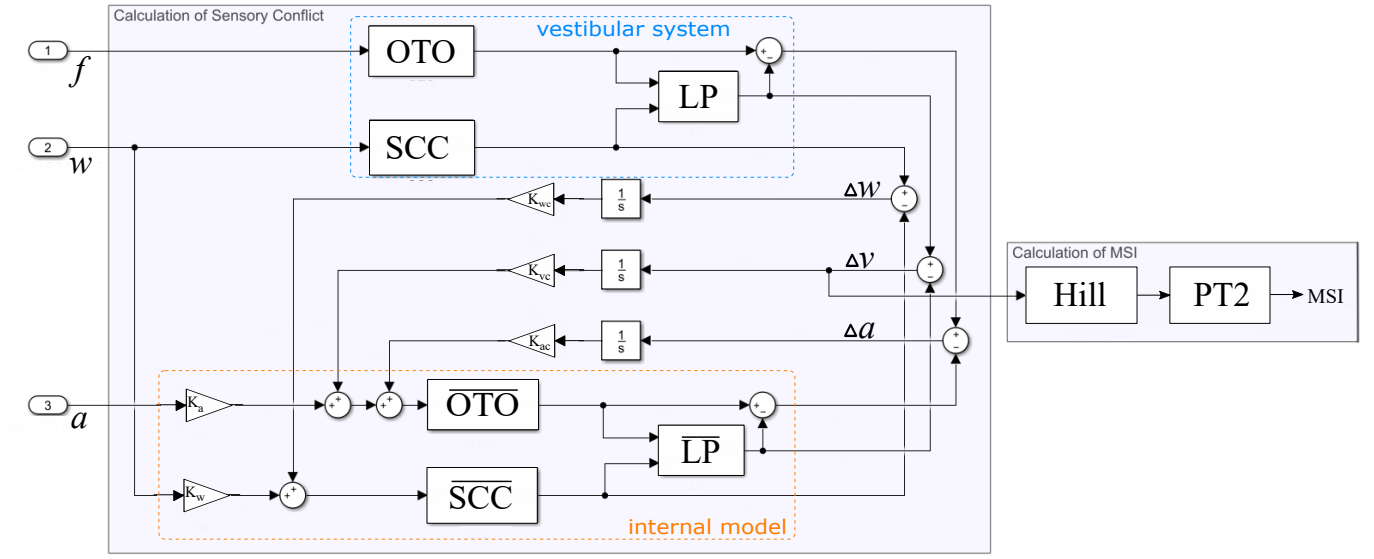
Original 6DOF-SVC model [28] applied on the five driving experiments. Model of the vestibular system framed in blue and internal physiological reference model framed in orange.

The 6DOF-SVC model [24], [34], [46], [57], see also [56], [58], estimates the SC by comparing sensory signals derived from the vestibular system, represented by models of the semicircular canals (*SCC*) and the otolith organs (*OTO*) [30], with corresponding signals generated by an internal reference model. Based on the estimated SC, MSI is subsequently computed over time. According to the definition of [32], [33], the MSI is equivalent to the incidence of vomiting. However, the model allows for an alternative definition, provided that the designated model parameters are used and the available data, or the training based on that data, supports such an approach. A detailed mathematical formulation is provided in the Appendix.

#### Input of the Model

The standard 6DOF-SVC model, Fig. 1, takes as input three-dimensional accelerations excluding gravity (*a*), three-dimensional accelerations including gravity (*f* ) and three-dimensional angular velocity (*w*), all referenced to a coordinate system centered at the passenger’s head. A multidimensional spatial and dynamic relationship exists between the vehicle reference point, represented by the location of the vehicle-mounted sensors, and the occupant’s head, which serves as the reference point for the model. This relationship is primarily influenced by two factors. First, a spatial offset is present between the two reference points, which is mainly determined by the seating position and the anthropometric characteristics of the occupant. Second, dynamic effects arise from active and passive occupant movements, including both head and body motions. In the case of body motion, different movement patterns and conditions can be distinguished, as extensively discussed in literature [24], [28], [57]. As previously investigated by the authors [29] the resulting MSI prediction errors are largely negligible when occupants perform NDRTs, as occupants typically remain in a quasi-static posture under a compliant head condition (representing a passenger without active MS counter body movements [28]). The offset between the passenger’s head and the position of the acceleration sensor of the vehicle can also be neglected, as the resulting error is minimal, on the order of 10^*−*3^, and thus negligible in relation to the absolute MSI values [29]. Furthermore, since the model parameters are calibrated specifically for the considered application scenario, inaccuracies in the representation of the underlying physical relationships may be partially compensated for during the parameter estimation process. Therefore, we used the driving dynamic profile, defined as [**f** (*t*) **w**(*t*) **a**(*t*)]^*T*^ in [m/s^2^, rad/s, m/s^2^], of the inertial measurement unit of the car, recorded for each measurement and each participant of the five experiments, as input to the 6DOF-SVC model. Moreover, natural variations resulting from manual vehicle operation, since no fully autonomous test vehicles were available, led to slight differences in driving dynamics across participants. Consequently, since several fluctuating driving dynamics are used as model input, each predicted MSI of every measurement also exhibits natural variability, see Fig. 2 (black lines). Therefore, the model validation process focused on a single, fully recorded driving dynamic profile per experiment, which represents the average driving dynamic profile on this test route, to predict the average MSI, see Fig. 2 (red line), which will be compared with the individual experimental MS occurrence.

**Figure 2:**
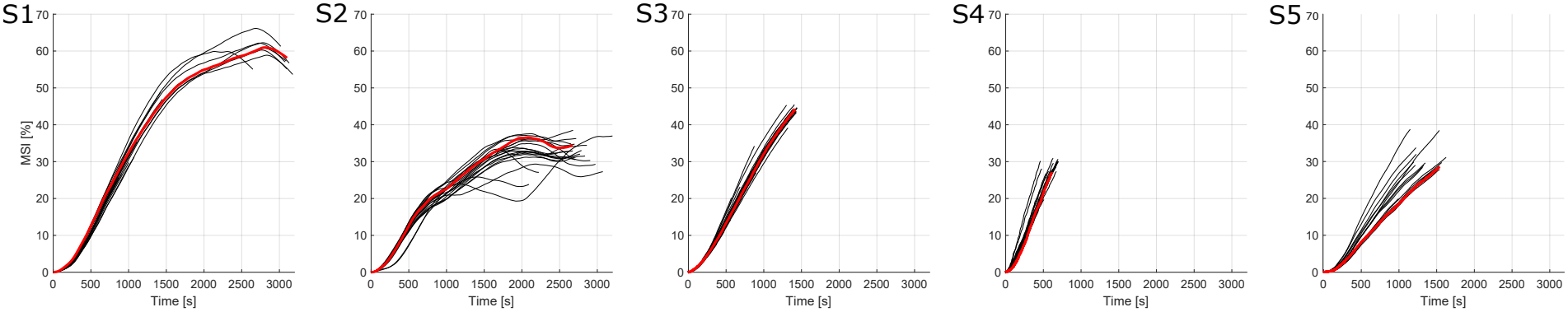
Predicted MSI of each measurement drive (each recorded driving dynamic profile) using parameter set *B*25. Reference (average) MSI colored in red.

### 2.3 Comparision between Prediction and Experimental Results

#### Experimental MSI

To compare the predicted average MSI of the model with the actual experimental results, an experimental MSI (expMSI) is determined based on subjective well-being of the participants of the experiment. As suggested by our previous work [29], we calculate the expMSI for each driving experiment (S1-S5) by determining the percentage of motion sick participants, who rated MISC of at least 7, throughout the temporal progression of the experiment. This definition of MS, using MISC of 7, is equivalent to the rating “fairly nausea” [44]. Furthermore, it can be regarded as a less restrictive extension of the original MSI definition [32], [33], which was primarily based on the probability of vomiting occurrence. To calculate the expMSI, every point in time when a participant reached an MISC rating of 7 for the first time (fairly nausea) is marked. Then, the percentage of participants who feel motion sick at these time points is determined. This results in the expMSI as a stepwise trajectory.

#### Prediction Performance Metric

To evaluate the performance of the prediction two metrics are used: First, the difference (Δ*MSI*) between the final predicted MSI value and the final expMSI value is determined. Second, to evaluate the accuracy of the predicted MSI trajectories, the Root Mean Square Error (RMSE) (eq. 1), between predicted MSI and the expMSI is calculated. Minimizing the RMSE ensures that the curve of the predicted MSI approximates the data points of expMSI as precisely as possible.

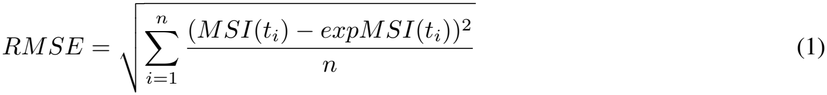

where *t*_*i*_ is the time point when a participant became motion sick (MISC *≥* 7); *n* represents the number of expMSI values (number of sick participants) of the single experiment.

#### Parameter Optimization and Validation

Two model parameter sets are suggested for MSI prediction in passenger groups that perform an NDRT in a real driving car; and were published previously by the authors [29]. Parameter set *B*20_1_, obtained through a conservative optimization of parameters *K*_*ac*_, *K*_*wc*_, *K*_*vc*_, *b*, and *P*, achieved high accuracy in predicting the final MSI value at the end of a drive. In contrast, parameter set *B*20_2_, resulting from a more comprehensive optimization of parameters *K*_*a*_, *K*_*w*_, *K*_*ac*_, *K*_*wc*_, *K*_*vc*_, *b, τ*, and *P*, provided substantially better performance in reproducing the complete MSI trajectory. With the availability of additional digital resources in recent years, the parameter sets will be re-optimized (parameter set *B*25). Therefore, related to our previous method [29], S1 and S2 are used as training data, and a Bayesian optimization algorithm with an expanded parameter range is employed, as shown in Table 2. For the analysis, Bayesian optimization with an exploration ratio of 0.7 and a maximum of 1000 objective evaluations was applied, implemented in MATLAB (R2019a, The MathWorks Inc.). For final optimization, the algorithm minimizes the sum of the RMSEs of S1 and S2 simultaneously, as shown in (eq. 2).

**Table 2:** Parameter Optimization Ranges.

| *K<sub>a</sub>* | *K<sub>w</sub>* | *K<sub>ac</sub>* | *K<sub>wc</sub>* | *K<sub>vc</sub>* |
|---|---|---|---|---|
| 0.0 - 5.0 | 0.0 - 2.0 | 0.0 - 3.0 | 0.0 - 10.0 | 1.0 - 10.0 |
| *b*[m/s<sup>2</sup>] | *P*[%] | *τ*[s] | *τ<sub>d</sub>*[s] | *τ<sub>a</sub>*[s] |
| 0.0 - 1.0 | 0.0 - 100.0 | 1 - 1200 | 7.0 | 190 |

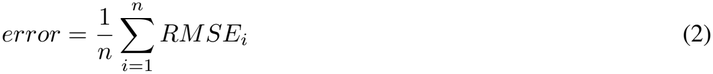

where *n* is set on 2 (i = experiment number)

Finally, see Table 3 for the parameter overview. Compared to *B*20_2_, *K*_*a*_, *K*_*v*_ and *K*_*ac*_ of *B*25 are similar to the previous values. *τ* is increased by 80 seconds, leading to a more stable MSI behavior. This is slightly compensated by an adjustment of *b*, which affects the increase of the predicted MSI. The optimized *P* value is slightly lower than that of *B*20_2_. The largest difference shows the strongly increased *K*_*vc*_ and *K*_*wc*_ values, which directly influence the duration and impact of the estimated SC.

**Table 3:** 6DOF-SVC Model Parameter Sets.

|  | *K*<sub>*a*</sub> | *K*<sub>*w*</sub> | *K*<sub>*ac*</sub> | *K*<sub>*wc*</sub> | *K*<sub>*vc*</sub> | *b*[m/s<sup>2</sup>] | *P*[%] | *τ*[s] | *τ*<sub>*d*</sub>[s] | *τ*<sub>*a*</sub>[s] |
|---|---|---|---|---|---|---|---|---|---|---|
| *B*20<sub>1</sub> | 0.1 | 0.8 | 0.35 | 0.8 | 1.0 | 0.1 | 88 | 720 | 7.0 | 190 |
| *B*20<sub>2</sub> | 4.28 | 0.74 | 0.013 | 3.67 | 3.98 | 0.81 | 98 | 417 | 7.0 | 190 |
| *B*25 | 4.81 | 0.66 | 0.014 | 8.14 | 9.09 | 0.47 | 95.5 | 495 | 7.0 | 190 |

### 2.4 Analysis of MSI Fluctuations on the Test Routes

Strong fluctuations in the driving dynamic profile can arise due to different driving styles (ranging from more passive to more aggressive behavior) or varying traffic conditions. Although the test routes of the experiments were closed to public traffic (in four out of five cases), and the human driver was trained and deliberately aimed to perform reproducible runs, natural variations still occurred due to environmental influences or human error. Consequently, the predicted MSI (model output) also fluctuates, see Fig. 2 as an example for parameter set *B*25, due to variations in the driving dynamic profiles (model input). In the second step of our analysis, we will therefore quantify the fluctuations in both model input and output with respect to the previously introduced reference measurements (average MSI) of the individual experiments. For this analysis, the model uses the *B*25 parameter set.

#### Driving Dynamic Profile Fluctuations (Input Side)

To enable a consistent comparison between the complex driving dynamics profiles (three dimensional acceleration and angular velocity), a simple and robust metric must be selected. However, no established method for input comparison currently exists for the 6DOF-SVC model. Therefore, we simplify the model input to an one-dimensional variation index, due to the limited number of real-world measurements. To compute this one-dimensional variation index, the mean and standard deviation of the vector magnitude of the three-dimensional acceleration (excluding gravity) are calculated first. Then, the normalized mean difference (normDist), representing the distance between both distributions, is determined using the mean and standard deviation of the reference measurement (eq. 3). If a driving dynamic profile is incomplete due to a participant aborting the measurement, the reference measurement is truncated at the corresponding termination point.

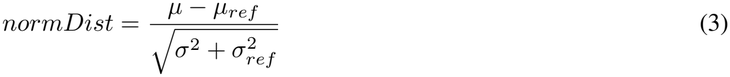

where *µ* and *σ* are the mean and standard deviation of the driving dynamic profile, and *µ*_*ref*_ and *σ*_*ref*_ are the mean and standard deviation of the reference driving dynamic profile.

#### MSI Fluctuation (Output Side)

Since the driving dynamics profiles exhibit temporal fluctuations, the MSI trajectory is disregarded, but both profiles reach the same route endpoint, so the final MSI value will be taken into account. Hereby, the percentage change/ deviation in MSI (pcMSI) is computed between the predicted MSI of a normal measurement and the reference, average MSI (eq. 4). As in the previous step, if a driving dynamic profile is incomplete due to a participant aborting the measurement, the MSI reference prediction is truncated at the corresponding termination point.

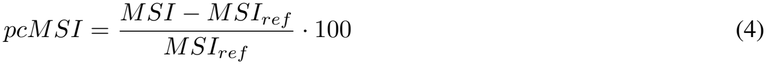

### 2.5 Calculation of an Individual MSI Threshold

To determine the individual MSI threshold, we identified the time points when participants became motion sick (MISC *≥* 7) and extracted the predicted MSI value (used model parameter set *B*25) for that moment, based on the participant’s driving session (individual driving dynamic profile). The overall dataset includes nine participants who took part in more than one study, as well as in the focused condition without CM. These cases are additionally processed to enable comparison across experiments.

### 2.6 Method Summary

For each of the five driving experiments, a single MSI is predicted using the three parameter sets (published *B*20_1_ and *B*20_2_ as well as optimized *B*25) and the driving dynamics, corresponding to the average driving dynamic profile of the test route, as input. This predicted “average” MSI is compared to the experimental MSI (expMSI), which was calculated based on the participants who felt motion sick (MISC *≥* 7; “fairly nausea”) in the experiment. The prediction error between the predicted and experimental MSI is quantified using Δ*MSI*, which evaluates the final MSI value, and RMSE, for evaluation of the MSI trajectory.

The second analysis step evaluates the MSI prediction deviation. Input side: For each recorded driving dynamics (each measurement) the normalized distance to the used “average” driving dynamic profile was calculated, see Fig. 3, to quantify driving dynamic fluctuations of the single driving experiments. Output side: On each recorded driving dynamic the MSI of each participant was predicted (using parameter set *B*25). Subsequently, the percentage deviation (pcMSI) of the predicted final MSI values is evaluated in comparison to the reference measurement of each experiment (“average” MSI).

**Figure 3:**
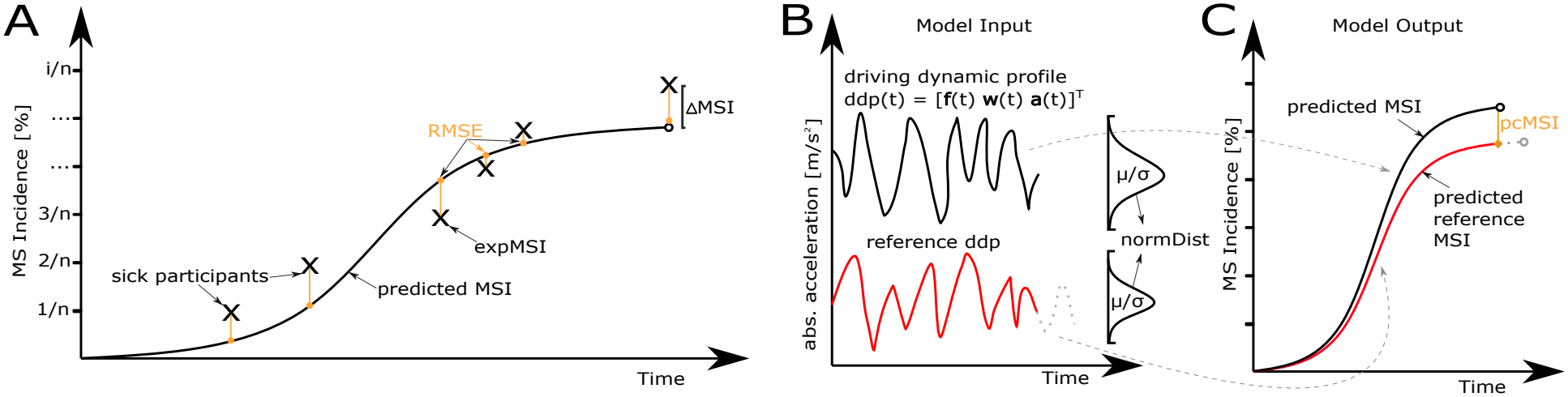
A) Comparison between predicted MSI of the 6DOF-SVC model (solid black line) and experimental results, i.e., motion sick participants of a single experiment (expMSI; x-marker). Prediction uncertainty quantified by the difference of the final value (Δ*MSI*) and the RMSE (orange) for the accuracy of the trajectory. B) Every measurement drive results in a driving dynamic profile (decribed by three-dimensional acceleration and angular velocity signals) used as model input. Deviations in the driving dynamic profiles are quantified by using a normalized mean difference (normDist). C) Every driving dynamic profile results in a predicted MSI. MSI based on the reference driving dynamic profile colored in red, representing the average drive of the driving experiment. Calculated percentage MSI deviation (pcMSI) to the reference drive colored in orange.

Final analysis step: The individual MSI thresholds at which participants experienced MS (MISC *≥* 7) are determined, based on the respective driving sessions (using parameter set *B*25).

## 3 Results

### Parameter Performance

The new optimized parameter set *B*25 was able to reduce the RMSE across most of the experiments (S1,S3-S5), see Table 4, resulting in a better predicted MSI trajectory compared to parameter set *B*20_2_. Moreover, unexpectedly, parameter set *B*25 also achieved the best prediction accuracy for the final MSI values of S3 and S4, see Table 5, thereby outperforming set *B*20_1_.

**Table 4:**
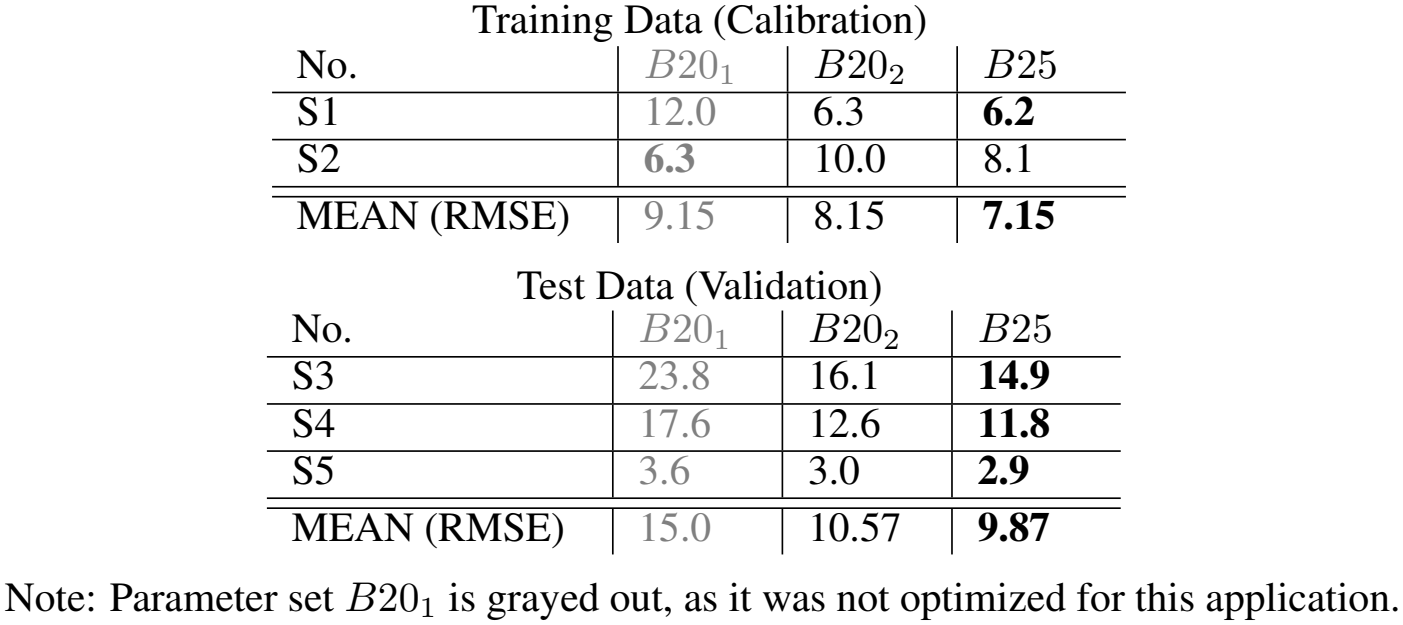
RMSE Values [percentage points] for the three different parameter sets. The parameters *B*20_1_ (gray) lie outside the intended scope of comparability but have been included for the sake of clarity and completeness.

| Training Data (Calibration) |  |  |  |
|---|---|---|---|
| No. | *B*20<sub>1</sub> | *B*20<sub>2</sub> | *B*25 |
| S1 | 12.0 | 6.3 | **6.2** |
| S2 | **6.3** | 10.0 | 8.1 |
| MEAN (RMSE) | 9.15 | 8.15 | **7.15** |
| Test Data (Validation) |  |  |  |
| No. | *B*20<sub>1</sub> | *B*20<sub>2</sub> | *B*25 |
| S3 | 23.8 | 16.1 | **14.9** |
| S4 | 17.6 | 12.6 | **11.8** |
| S5 | 3.6 | 3.0 | **2.9** |
| MEAN (RMSE) | 15.0 | 10.57 | **9.87** |

Note: Parameter set *B*20<sub>1</sub> is grayed out, as it was not optimized for this application.

**Table 5:**
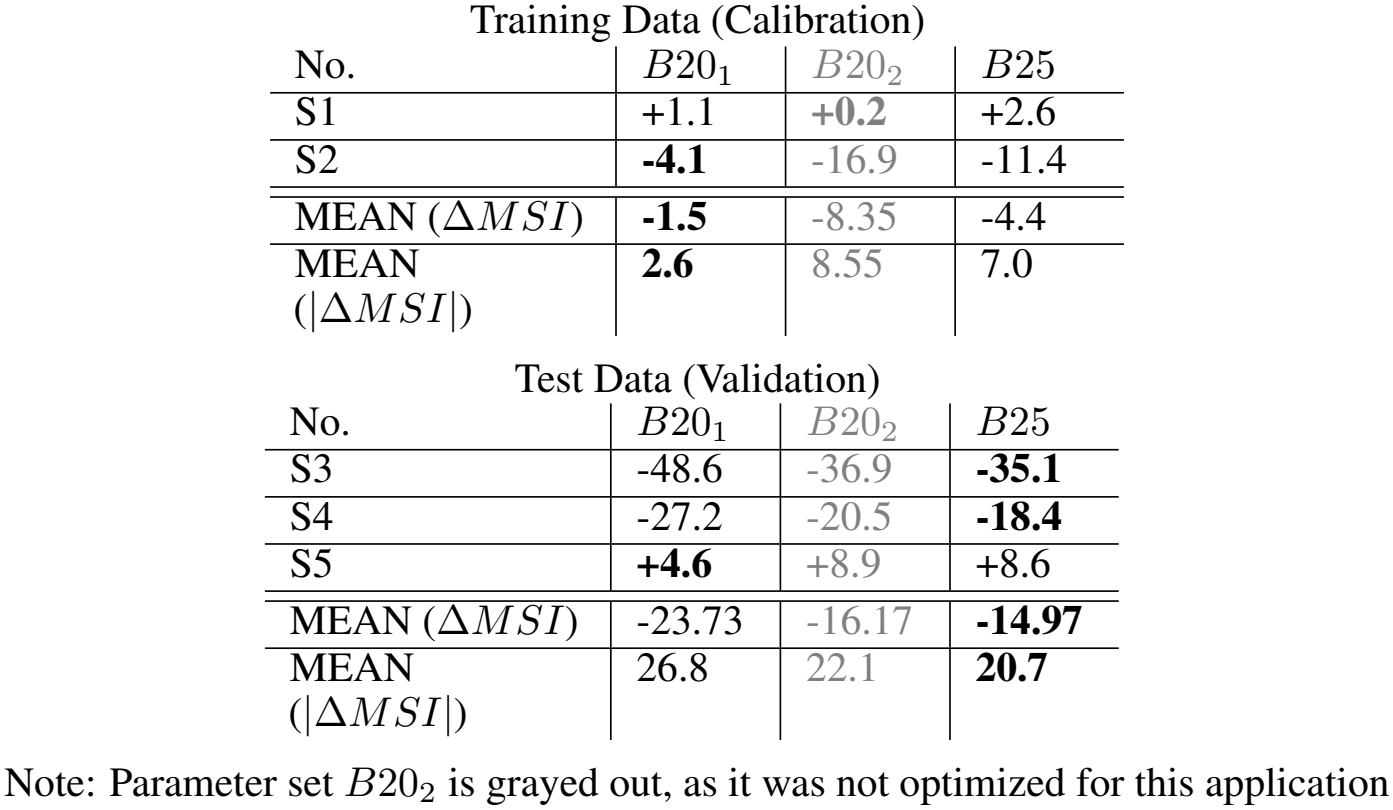
Difference of final MSI value (Δ*MSI* [percentage points]) for the three different parameter sets. The parameters *B*20_2_ (gray) lie outside the intended scope of comparability but have been included for the sake of clarity and completeness.

| Training Data (Calibration) |  |  |  |
| No. | *B*20<sub>1</sub> | *B*20<sub>2</sub> | *B*25 |
|---|---|---|---|
| S1 | +1.1 | **+0.2** | +2.6 |
| S2 | **-4.1** | -16.9 | -11.4 |
| MEAN (Δ*MSI*) | **-1.5** | -8.35 | -4.4 |
| MEAN (\|Δ*MSI*\|) | **2.6** | 8.55 | 7.0 |
| Test Data (Validation) |  |  |  |
| No. | *B*20<sub>1</sub> | *B*20<sub>2</sub> | *B*25 |
| S3 | -48.6 | -36.9 | **-35.1** |
| S4 | -27.2 | -20.5 | **-18.4** |
| S5 | **+4.6** | +8.9 | +8.6 |
| MEAN (Δ*MSI*) | -23.73 | -16.17 | **-14.97** |
| MEAN (\|Δ*MSI*\|) | 26.8 | 22.1 | **20.7** |

Note: Parameter set *B*20<sub>2</sub> is grayed out, as it was not optimized for this application

### MSI Trajectories

The pronounced increase of MS in S4 around 550 seconds was not predicted, resulting in an underestimation of the final MSI value in all three predictions, see Fig. 4 (d). Similarly, the sharp increase at the end of S3 could not be predicted; however, the MSI trajectory generally captures the expMSI, see Fig. 4 (c). S3 is predicted with a significantly higher MSI than S4 and S5, which is consistent with the expMSI.

**Figure 4:**
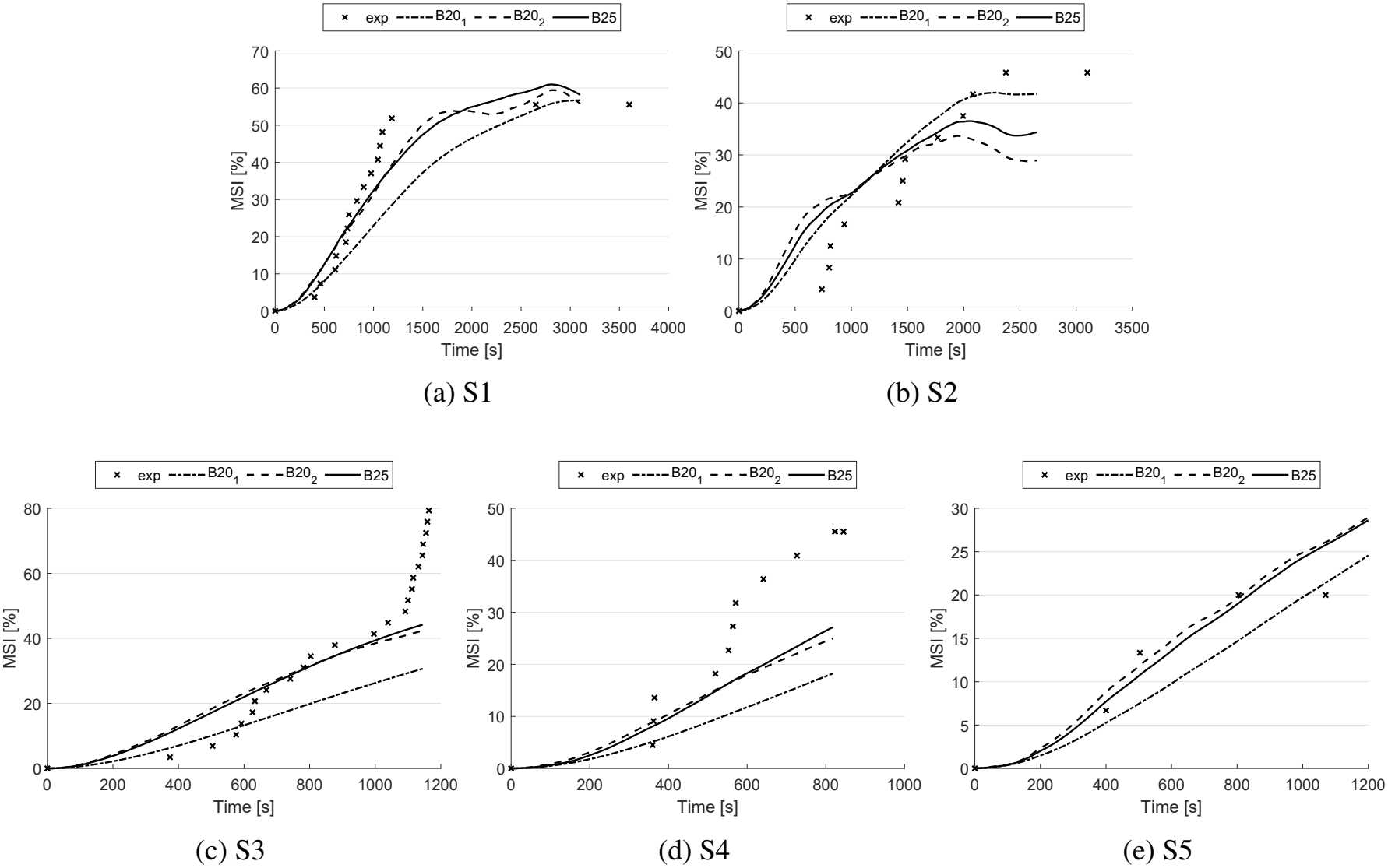
Predicted MSI of the five experiments compared to experimental results.

### MSI Fluctuations

The experimental results demonstrate a positive correlation (R = 0.59, p *<* 0.001) between normDist of the driving dynamics profile and the resulting MSI deviation, as expected (see Table 6 and Fig. 5; trendline: *pcMSI* = 101.487 *· normDist* + 0.087). Variations in the driving dynamics profile across individual measurements lead to a standard deviation in the final MSI value ranging from approximately 2% (in highly reproducible test drives, e.g., S4.1 and S4.2) to around 16% (in less reproducible drives, e.g., S5), as shown in Table 6. S2, the only experiment conducted in real-world traffic, exhibits a standard deviation of 9.71%. An analysis of all data points reveals an overall standard deviation of 10.18% (*σ*_*pcMSI*_ ). S4 illustrates two clearly separated clusters corresponding to different measurement sessions (week 1 and week 2), indicating an bias from the driver. Since the second cluster (S4.2) is not centered around zero, see Appendix Fig. 7 E, it was excluded from the evaluation. The data suggest that these two clusters correspond to the two different measurement weeks or the trial order of the experiment. In the second week/ trial, the driving dynamics profile is noticeably lower, yet the MSI level is approximately 12% higher, see Fig. 5.

**Table 6:** Percentage changes/ deviation in MSI (pcMSI) [%] for the different experiments.

| No. | pcMSI: mean (std) | Pearson’s r |
|---|---|---|
| S1 | -2.60 (3.97) | 0.72 |
| S2 | -7.52 (9.71) | 0.74 |
| S3 | 1.05 (4.83) | 0.87 |
| S4 | 6.57 (6.22) | -0.90 |
| S4.1 | 0.55 (2.18) | -0.01 |
| S4.2 | 12.03 (2.02) | 0.06 |
| S5 | 11.39 (15.89) | 0.33 |
| MEAN | 2.48 (6.43) |  |

**Figure 5:**
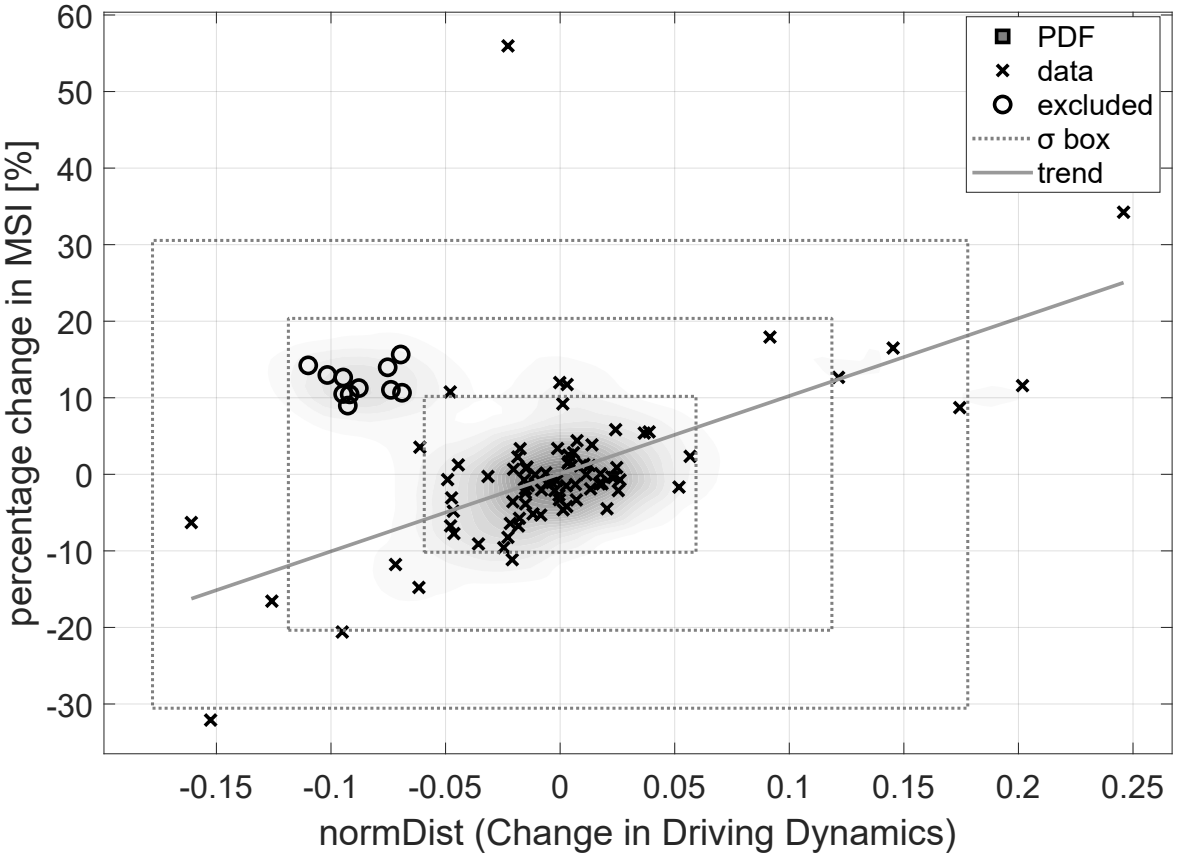
Relation between changed driving dynamic profile (as normalized mean difference (normDist) to the reference driving dynamic profile of the experiment) and percentage change in predicted MSI over all participants (R = 0.59, p *<* 0.001). Probability Density Function (PDF) colored in gray-level shading and framed by dotted, zero-centered 1, 2 and 3-*σ* boxes of the data (*σ*_*x*_ = 0.059 / *σ*_*y*_ = 10.18%). Trend line colored in light gray with the formula: *pcMSI* = 101.487 *· normDist* + 0.087. S4.2 cluster is excluded for trend and sigma calculation, because it is not zero-centered.

### Individual MSI Threshold

The experimental results demonstrate a negative, but not significant, correlation between the Q-Score and the individual MSI thresholds, see Fig. 6 and Table 7. S5 exhibits clear outlier behavior; however, it should be noted that only three data points are available for this experiment. In contrast, S1-S4 show a remarkably consistent trend, characterized by a similar gradient. However, the results of S4 are generally shifted toward lower MSI threshold values, both at the overall experiment level (Fig. 6) and on an individual participant basis (Table 7).

**Table 7:** MSI threshold (defined as MSI at the time point where MISC *≥* 7) for participants completing multiple experiments (S2-S5).

|  | MSI threshold [%] (Q-Score/max.MISC) |  |  |  |  |
|---|---|---|---|---|---|
| Subject-No. | S2 | S3 | S4 | S5 | Range |
| 1 | 31.47 (8/7) | 23.80 (10/10) | 16.60 (11/9) | 31.10 (12/8) | 14.87 (4/3) |
| 2 | NS (6/5) | NS (6/4) | 27.93(5/7) | - | - (1/3) |
| 3 | - | 41.54 (5/8) | 19.69 (6/10) | - | 21.85 (1/2) |
| 4 | - | 40.64 (6/7) | NS (7/4) | - | - (1/3) |
| 5 | - | 40.60 (11/9) | 26.88 (9/7) | - | 13.72 (2/2) |
| 6 | - | 25.33 (9/10) | 14.03 (8/9) | - | 11.30 (1/1) |
| 7 | - | NS (8/4) | NS (9/2) | - | - (1/2) |
| 8 | 19.92 (10/10) | 27.63 (11/10) | - | NS (11/5) | 7.71 (1/5) |
| 9 | 33.35 (9/9) | - | 18.43 (10/8) | 38.75 (10/7) | 20.32 (1/2) |
| MEAN |  |  |  |  | 14.96 (1.44/2.56) |

- : no participation; NS: not sick

**Figure 6:**
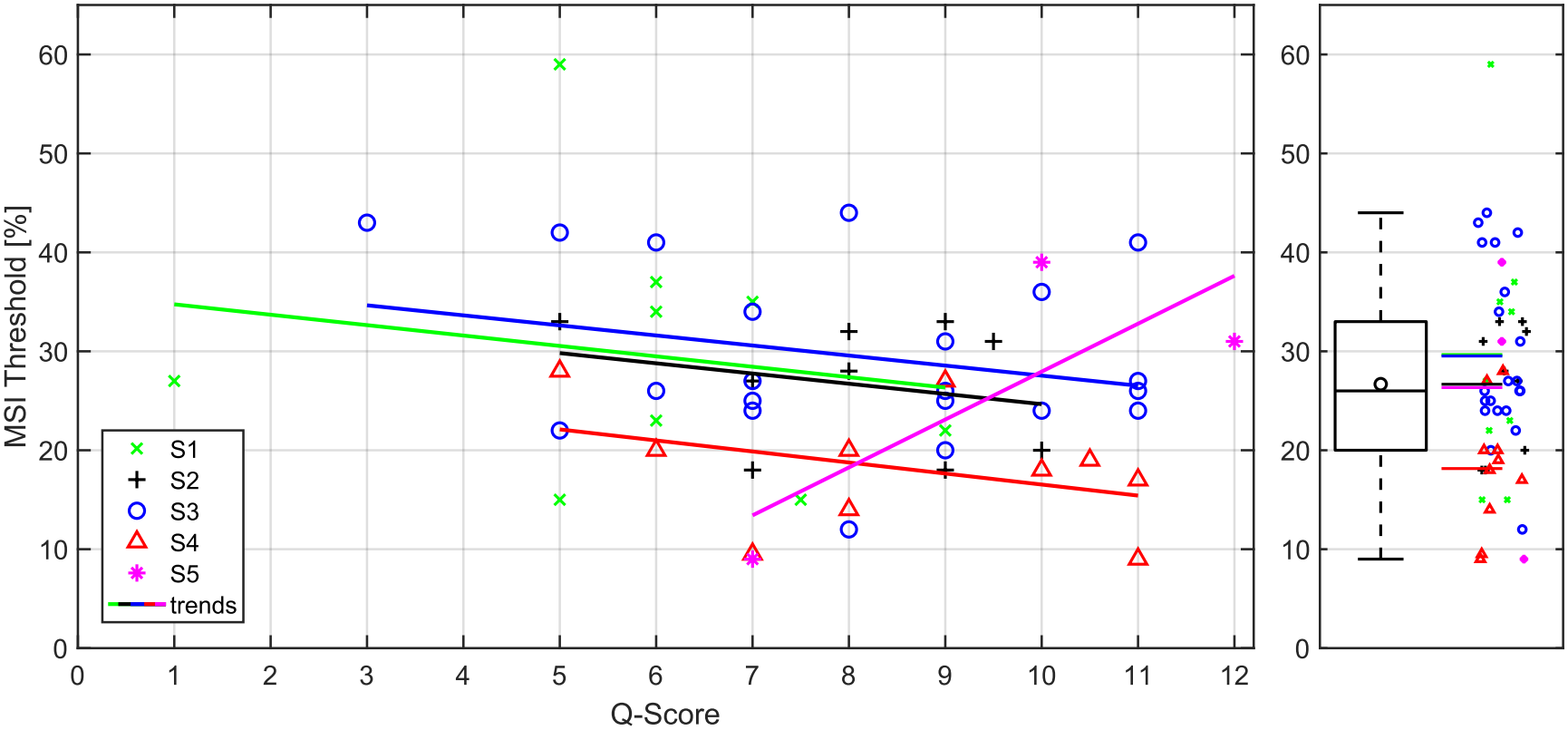
MSI thresholds at which individual participants first reported MS were plotted against the Q-Score, with data points annotated according to the corresponding driving experiment (S1-S5).

**Figure 7:**
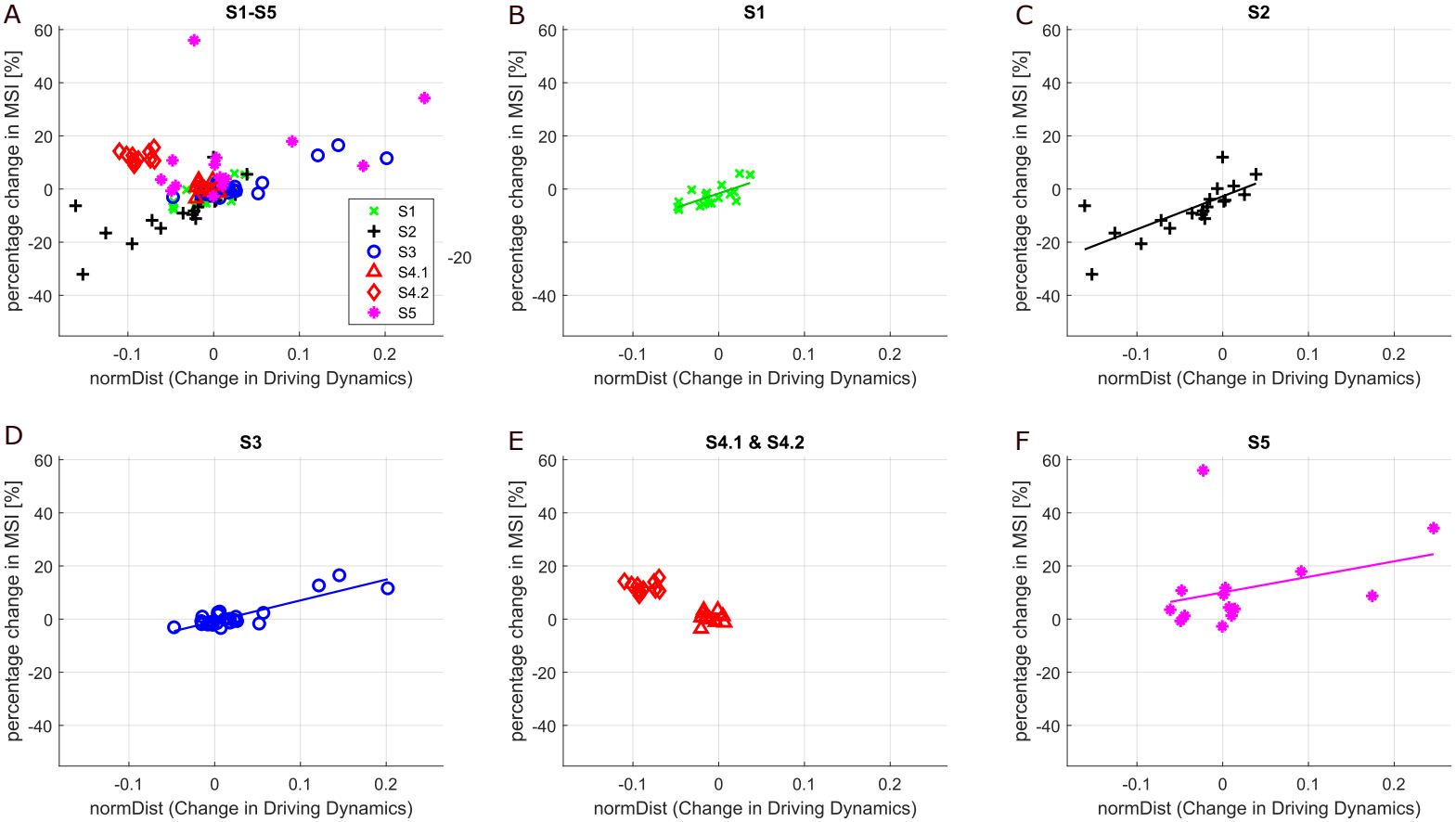
Relation between changed driving dynamic profile (as normalized mean distance (normDist) to the reference driving dynamic profile of the experiment) and percentage change in predicted MSI for each experiment. Experiment 4 was divided into two subgroups based on the emergence of two distinct clusters (S4.1 and S4.2). Furthermore, the trend curves have been sketched within the relevant data range.

**Figure 8:**
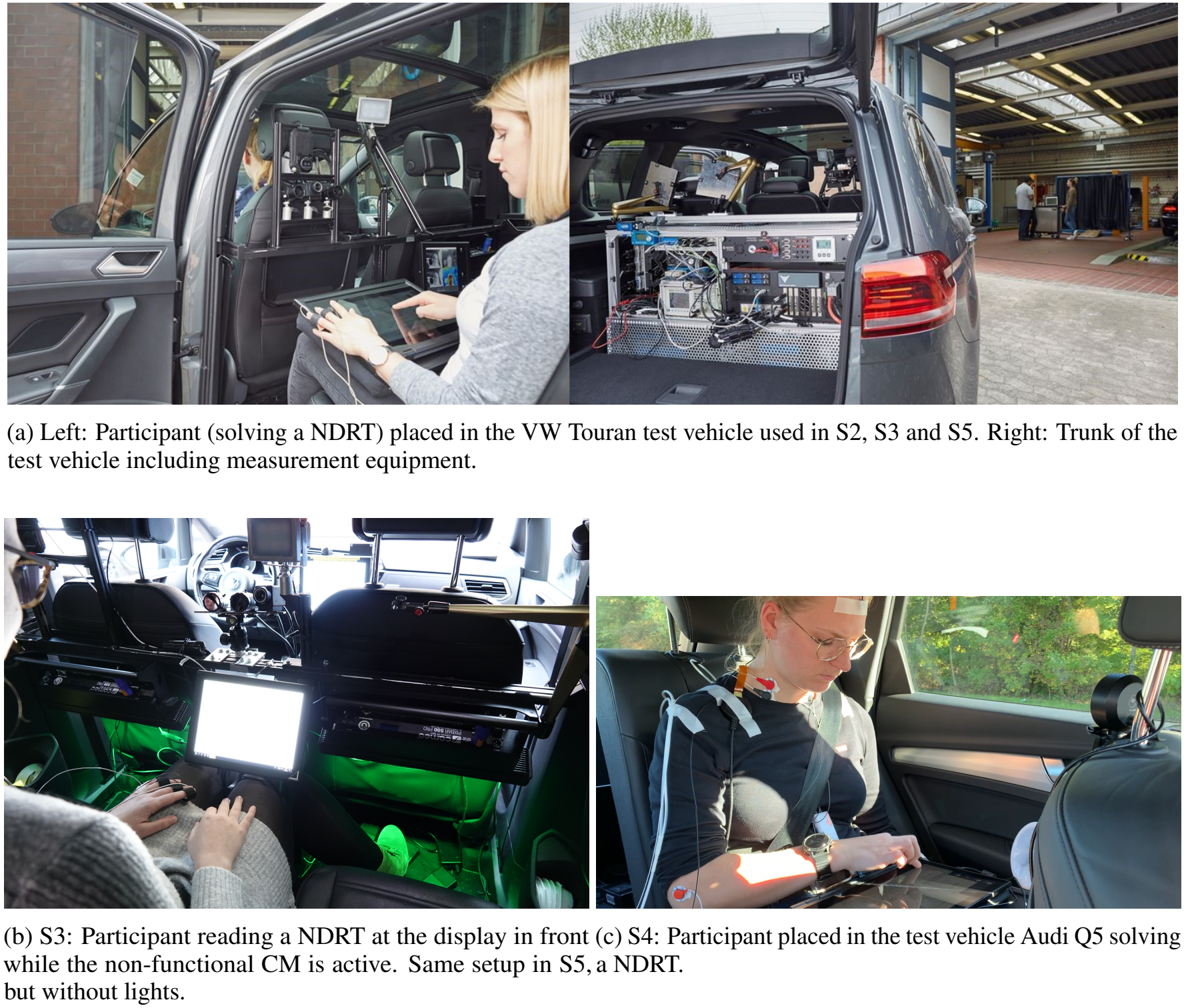
Experimental setups.

## 4 Discussion

### 4.1 Parameter Performance

Based on the higher prediction accuracy of both performance parameters, MSI trajectory and final MSI value, see RMSE and Δ*MSI*, Table 4 and 5, we recommend discontinuing the use of both older parameter sets in favor of the newly developed parameter set *B*25 to predict MSI in passenger groups solving a NDRT. The new parameter set is characterized by a slightly more dynamic response of the model to SC, achieved through adjustments to parameters such as *K*_*vc*_ and *K*_*wc*_, see Table 3. This increased responsiveness is compensated by a temporal stabilization of the MSI trajectory via the time constant *τ* of the PT2 element, in comparison to *B*20_2_. The prediction accuracy for S3 and S4 was slightly improved (regardless of the absolute prediction error), which may indicate that further optimization iterations could enhance the predictive performance of the 6DOF-SVC model and that its full potential has not yet been reached.

### 4.2 MSI Prediction Accuracy

Despite further parameter optimization based on S1 and S2, the model is unable to closely reproduce the expMSI trajectories, see Fig. 4 (a)-(b). Instead of interpolating between the experimental MSI observations, the model provides only an approximation of the measured trajectories. The remaining discrepancies indicate that the limitations are not primarily caused by suboptimal parameter values, but rather by structural simplifications of the model formulation itself, particularly the employed Hill function and PT2 element, which is consistent with previous findings [29] as well as the trend to replace them with an Oman-Nausea-Submodel [37], [46], [47]. Consequently, even after calibration, the model is unable to perfectly align with the experimental MSI data. While this imperfect fit may initially appear disadvantageous, it also suggests that the model is not merely overfitting the calibration datasets. The persistence of prediction errors, even for the datasets used during parameter optimization (S1 and S2), indicates that the model retains a certain degree of independence from the calibration data. This behavior can be interpreted as an indication of model robustness, as the resulting MSI predictions are not excessively tuned to individual datasets. S3 exhibits the lowest prediction accuracy, with MSI trajectories diverging extremely after 1000 seconds from the start of the experiment. But, S3 is based on a truncated distributed population as there are more susceptible participants, see elevated Q-Score in Table 1. This likely leads to an increased expMSI, which the 6DOF-SVC model cannot accurately predict, as it was trained on a homogeneous participant group (S1 and S2). From a purely technical and application oriented standpoint, the 6DOF-SVC model should be capable of predicting the elevated expMSI observed in S3 by increasing the value of *P*, potentially even beyond 100%. Assuming our predicted MSI of S3 is accurate and valid for a homogeneous passenger group, it can be concluded that fluctuations in MS susceptibility within the target population may lead to deviations in MSI of up to 35 percentage points, see Table 6. This highlights the importance of specifying the initial MS susceptibility of the study population in experiments, not only for further model development but also for accurately interpreting study results and meta-analyses.

The 6DOF-SVC model appears to struggle with predicting the characteristic contours of expMSI trajectories, regardless of the parameter set used. The model still fails to predict the high expMSI of S4 and the lower ones for S2, as shown in Fig. 4. Technically, the model is capable of exhibiting significantly more unstable behavior through the parameter *b*, thereby allowing for the representation of dynamic MSI patterns. However, optimization algorithms tend to converge towards solutions that favor sluggish or inert MSI dynamics. S4 focused on many lateral maneuvers, e.g., slalom driving without significant vertical load. S2 was driven in real traffic without designed lateral maneuvers, but including some speed bumps. It is possible that the 6DOF-SVC model is unable to sufficiently learn this distinction, as the *K*-parameters are uniformly applied across all three spatial input dimensions, resulting the SC is not weighted in any of the three directions. Although vertical acceleration is known to be a strong trigger for MS [31], [32], horizontal acceleration within the vehicle, particularly under the influence of a NDRT and the resulting altered head orientation relative to the vehicle, has not yet been sufficiently investigated. Consequently, it remains unclear how the SC manifests across the three spatial dimensions. This uncertainty also raises the question of whether the model should deviate from the generalization of the SC or the *K*-parameters. However, it must be considered that introducing additional parameters may lead to increased model complexity, potential overfitting, and reduced generalization and stability.

The predicted MSI trajectories show significant deviations both before and after optimization (Fig. 4). Characteristic plateaus and slopes observed in the expMSIs trajectories are not reproduced by the predicted MSI of the model, which, as discussed previously, indicates that the PT2 element is not an optimal post-processing of the SC. One of the most notable discrepancies in the MSI trajectories is the expMSI onset, particularly evident in experiment S2. S2 was conducted in real-world traffic conditions and likely represents one of the mildest driving dynamics profile compared to the other test tracks of S1, S3-5, see Appendix Fig. 10. Despite S2 being part of the training dataset, the 6DOF-SVC model fails to reproduce this temporal offset. This limitation suggests that the model may not adequately capture the delayed onset of MS under smoother, real-world driving conditions. MS develops over time because the body must be exposed to stimulation before symptoms occur. The five experiments indicate that there is a delay of about 5 minutes before the first participant experiences severe nausea. Mathematically, however, an SC is calculated during this time, which increases the MSI just a few seconds later. Because this delay also varies across individuals and results in the expMSI, it has to be further investigated how this time shift in MS onset can be included in the 6DOF-SVC model.

### 4.3 MSI Fluctuations

The experimental results show that the normDist of the driving dynamics profile positively correlates with the resulting MSI deviation, reflecting the technical behavior of the 6DOF-SVC model. Therefore, the normDist can be considered as an useful one-dimensional reference metric for evaluating the multidimensional model input of the driving dynamics profile. But, the cluster offset between S4.1 and S4.2 represents a limitation. Despite a lower normDist in S4.2, the MSI was significantly higher. This behavior contradicts the trendline and can be explained by the assumption that, during this week, the test driver drove less aggressively overall but likely hit a slalom frequency closer to 0.2 Hz, known as strong trigger for MS [32]. The normDist does not capture frequency-specific effects and was computed solely based on acceleration, excluding angular velocity. Although the results are generally positive, this relationship highlights a limitation that must be acknowledged, as it may contribute to the variability observed in the data, Fig. 5. This deviation also demonstrates that even when the same route is driven with the same intention (i.e., reproducibility), an MSI deviation of up to 11.48% (Δ*MSI*_*S*4.2_ *−*Δ*MSI*_*S*4.1_; Table 6) can occur. Considering the standard deviations of S2 (experiment in real-world traffic) of 9.71%, it can be assumed that both the predicted and experimentally measured MSI for a given route may fluctuate by approximately *±*10% only due to variations in driving dynamics.

This insight is fundamental for the application of the 6DOF-SVC model. First, it suggests that the derivation/ error of the predicted MSI trajectory likely falls within an acceptable error margin. Second, it suggests that the potential for MSI reduction through targeted adjustments to driving strategy (e.g., passive vs. aggressive driving) is likely greater than 10%. In future vehicle applications, this could enable comparisons of MSI between an alternative route and a different driving style on the same route. Our trendline can serve as a new tool for intelligent vehicles to estimate the benefit of a more passive driving style. However, this should be validated through further targeted studies on driving behavior. Third, given the magnitude of this uncertainty, it is worth questioning whether MSI, both predicted and experimentally measured, can serve as a reliable indicator of MS. The error may be too large to develop a stable parameter set with high prediction accuracy. Therefore, more driving experiments have to be conducted in highly controlled environments (e.g., autonomous vehicles or driving simulators) to ensure greater reproducibility of driving dynamics. Nonetheless, the development of MSI databases should continue to promote the development of reliable MSI models.

### 4.4 Individual MSI Threshold

Excluding S5 due to the availability of only three data points, the remaining four experiments exhibit a negative correlation between MSI threshold and Q-Score (Fig. 6 and Table 8). This observation is consistent with the expectation that participants with a higher susceptibility to motion sickness, as identified by the Q-Score questionnaire, experience the onset of nausea at lower MSI thresholds. Nevertheless, the variability in MSI thresholds is sufficiently large that the observed trend is not statistically significant. Overall, the MSI thresholds appear approximately normally distributed around 26.69%. Considering a standard deviation of 10.04%, the critical range for actual nausea during driving is estimated between a predicted MSI of 16.65% and 36.73%. This additional information could support the application of the 6DOF-SVC model in navigation systems or route planning, for instance, by planning rest areas or fuel stops. The observation that participants of S4 exhibit lower MSI thresholds (Table 7 and Fig. 6) may be related to the generally underestimated MSI predictions for S4, which show high prediction errors (RMSE and final MSI; see Table 4 and 5). Considering the general MSI offset, the MSI threshold ranges, averaging 14.96 percentage points, may be narrower for individual participants than the small sample size suggests, see Table 7. Nevertheless, even with an individualized MSI threshold for specific car passengers, deviations from the norm must be assumed, as daily fluctuations could lead to increased or decreased susceptibility to MSI during the ride, which cannot be ruled out. In summary, the results suggest that MSI thresholds are influenced by an individual’s baseline susceptibility to MS. However, the observed variability appears too large to support reliable application in their current form. This conclusion is further supported by the fact that the MSI thresholds of individual participants vary considerably across different experiments. Nevertheless, an important finding of this study is that the identified MSI thresholds may enable the localization of critical periods along a route during which occupants are most likely to report the onset of nausea when using this modeling approach. Consequently, even if MSI thresholds are not sufficiently robust for direct

**Table 8:**
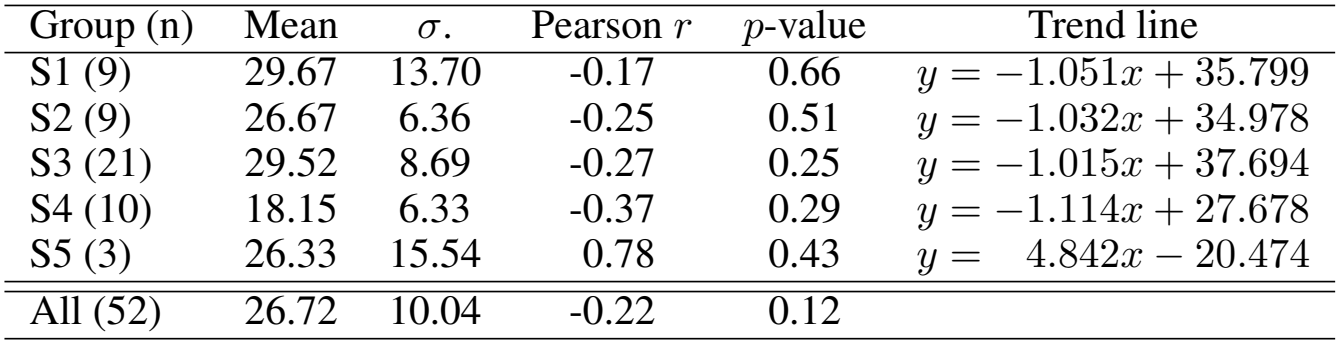
MSI threshold statistics, Pearson correlation coefficients, and corresponding linear trend lines for the individual driving experiments (S1-S5) shown in Fig. 6.

| Group (n) | Mean | *σ*. | Pearson *r* | *p*-value | Trend line |
|---|---|---|---|---|---|
| S1 (9) | 29.67 | 13.70 | -0.17 | 0.66 | *y* = −1.051*x* + 35.799 |
| S2 (9) | 26.67 | 6.36 | -0.25 | 0.51 | *y* = −1.032*x* + 34.978 |
| S3 (21) | 29.52 | 8.69 | -0.27 | 0.25 | *y* = −1.015*x* + 37.694 |
| S4 (10) | 18.15 | 6.33 | -0.37 | 0.29 | *y* = −1.114*x* + 27.678 |
| S5 (3) | 26.33 | 15.54 | 0.78 | 0.43 | *y* = 4.842*x* − 20.474 |
| All (52) | 26.72 | 10.04 | -0.22 | 0.12 |  |

**Table 9:**
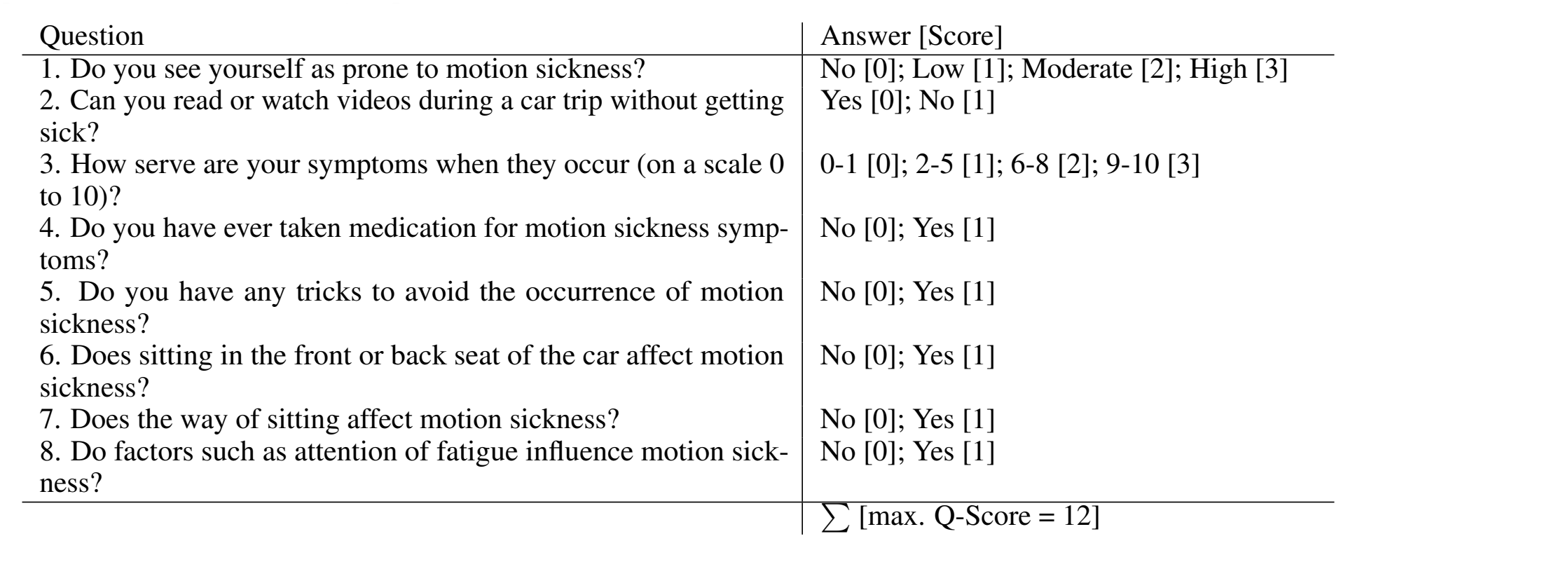
Pre-experiment, 8-item MS susceptibility questionnaire [54] to calculate the individual Q-Scores of the participants. The questionnaire comprises items inspired by established indicators and predictors of MS susceptibility, including self-assessed susceptibility [44], [59], the use of MS medication [60], and individual coping or avoidance strategies [61]. Additional items addressed the influence of seating position [16], [61], fatigue [62], mental distraction and attentional factors [7], as well as engagement in NDRTs [63]. To minimize potential placebo effects and negative expectancy biases toward the experiment, which have previously been shown to influence MS outcomes [20], the questionnaire was intentionally kept concise and non-intrusive.

### 4.5 Assessment of Model Applicability

Although the predicted MSI trajectory deviated from the experimental results, the trend of MS occurrence in the experiments was nevertheless correctly reflected in the predicted MSIs. So, the 6DOF-SVC model (using set *B*25) predicts the MSI trajectories of S3 and S4 to be higher than those of S1, and at the same time, S5 is similar to S2. This represents the correct trend of the three additional experimental results (S3-S5) in relation to the training data (S1, S2). Based on our previous findings, an MSI inaccuracy of *±*10% cannot be ruled out (see MSI fluctuations). Accordingly, all five predicted MSIs can be considered reasonably plausible. As a result, the 6DOF-SVC model indicates whether a route is likely to induce a high or low MSI, usable in navigation systems or route planning algorithms to choose the best driving route for MS susceptible passengers. If the accuracy of the MSI trajectory can be further improved in the future, it may enable a smart vehicle to determine at which point on the route a predefined critical MSI value is reached, e.g., to assess which MS avoidance strategies (e.g., alternative driving route or technological CM) would be helpful for the passenger. Mapping the predicted MSI to an individual passenger’s personal MSI threshold currently appears to have limited practical applicability, as both the expected inaccuracies in MSI prediction and the pronounced variability of personal MSI thresholds across multiple trips result in excessive overall deviations. For a conclusive assessment of individual MSI thresholds, additional multi-session driving experiments with the same participants should be conducted, preferably including longer and less intense measurement drives (e.g., extended highway trips rather than closed, purpose-built test maneuvers). Consequently, whereas the 6DOF-SVC model demonstrates clear potential for on-board applications, the development of a large-scale training database, ideally based on real-world MS driving experiments, likely represents the next essential step toward making the model truly ready for practical deployment.

### 4.6 Limitations

The reproducibility of MS experiments in real vehicles remains unclear, primarily due to the influence of traffic and individual differences. There are different individual aspects, like time of sleep, food intake, hormonal changes, and psychological factors, which precise quantitative influence on MSI values remains unclear and which are difficult to control in experimental settings. It should also be noted that the MSI used in this study represents severe symptoms (MISC *≥*7). Although it is plausible that the proposed concept may also apply to milder symptoms, this has not yet been demonstrated. In addition, expMSI is based on subjective data and is influenced by the interval of the MISC queries. Therefore, it is possible that the expMSI inherently contains a methodological offset, which may contribute to the observed temporal discrepancies. Additionally, a different perception of the initial symptoms may be observed across individuals.

Furthermore, the MSI prediction is subject to the error that the body and head movements of the passengers as well as the offset between head and sensor placement are simplified. Even if the error is expected to be very small based on previous findings [29], and may be partially absorbed by the model calibration process, it would be advisable to record the body movement of the participants to ensure that the assumptions regarding occupant motion within the 6DOF-SVC model are satisfied. Alternatively, the accelerations and angular velocity for the input of the model could also be measured directly at the passenger’s head via an external measurement unit. It should further be noted that the transfer of findings between the individual studies relies on the assumption of a high degree of similarity between the experimental conditions. This assumption represents a limitation of the present analysis and may affect the generalizability of the reported model performance. Across the five experiments, three different test vehicles were used. Although these vehicles are similar in type, variations in their driving dynamics profiles have to be expected. Such differences may also influence the expMSI and MSI prediction accuracy, introducing an additional source of variability that should be taken into account when interpreting the results. In particular, S2, S3, and S5 were conducted using the same vehicle platform, whereas S1 employed a comparable family van and S4 was conducted in an SUV. As detailed information regarding vehicle dynamics, modal characteristics, and direct cross-vehicle comparisons is not available, vehicle-specific effects cannot be excluded and may have contributed to the observed differences between the studies. Similarly, differences in seating position may introduce additional variability. The seating position was identical in S1, S2, and S4 (rear-right seat), whereas participants were seated in the rear-center position in S3 and S5. While the rear-center position reduces the lateral offset between the occupant and the vehicle centerline, the influence of seating position cannot be completely excluded. Differences in NDRTs and associated visual stimuli constitute another source of uncertainty. These effects cannot be explicitly represented within the current model structure and are therefore only indirectly reflected through the calibrated model parameters. Since validated models capable of consistently accounting for mixed NDRT conditions are currently lacking, the influence of NDRT-specific effects remains an open research question. Another potential offset arises from the fact that the average MSI determined in this study is based on the average driving profile, which may not necessarily correspond to the actual overall average of the test tracks. If additional data were available, the model could possibly increase prediction accuracy. Four of the five experiments were conducted in a closed testing area with specially designed intense maneuvers, which increase the MSI compared to experiments conducted in real road traffic. For realistic applications of the model, more data recorded in real traffic should be used to validate the 6DOF-SVC model. At the same time, it must be acknowledged that only five expMSI datasets were available for this investigation, which constitutes a significant statistical limitation. Publicly available datasets containing synchronized vehicle dynamics and experimentally determined MSI trajectories are currently not available. Consequently, additional datasets would be highly valuable for future research, particularly to assess whether the observed stability and plausibility of the model predictions persist across a larger number of experiments, vehicles, participants, and operating conditions. In addition, the model version employed in this study is exclusively vestibular-based, which may generally exhibit inferior performance compared to more recent models that integrate visual inputs. Moreover, ongoing research on observer models, as well as their combination with SVC models, appears poised to supersede the parameter adjustment strategy employed in this study, which is prone to unintended side effects.

## 5 Conclusion

Based on five real-world driving studies, which induced motion sickness in passengers (n = 105) who performed a non-driving-related task, two established parameter sets of the 6DOF-SVC model were outperformed by a newly optimized parameter set in terms of both MSI trajectory and final MSI values. Validation of the optimized parameter set demonstrates that previously unknown driving routes can be classified realistically with respect to their resulting MSI levels. However, notable discrepancies between the predicted and the experimentally observed MSI trajectories persist. Using the optimized parameter set in combination with the five available driving experiments, it was shown for the first time that MSI predictions can vary by approximately 10% solely due to natural fluctuations in driving dynamics, even under deliberately controlled and reproducible driving styles. This finding suggests that targeted adaptations of driving behavior, such as a consistently more passive driving style, could realistically lead to a reduction in motion sickness incidence exceeding 10%. Furthermore, the analysis of individual MSI thresholds at which participants actually experienced MS indicates that the area of interest for route planning should lie within an MSI range of 16.65% to 36.73%. However, the individual MSI thresholds exhibit substantial variability across different driving scenarios, which considerably limits their practical applicability and supports the alternative approach of predicting the MISC using the 6DOF-SVC model.

## 6 Acknowledgment

The authors want to thank Florian Dauth, Sven Greger as well as ZF Friedrichshafen AG for partially funding and providing the research vehicle as well as for technical support during the measurements. Furthermore we want to thank the Federal Ministry of Education and Research of Germany for funding of this research project [grant number: BMBF-FZ 13FH737IX6 and BMBF-FZ 13FH169KX0].

During the preparation of this work, the author used Microsoft Copilot in order to improve the readability and language of the manuscript. After using this service, the author reviewed and edited the content as needed and takes full responsibility for the content of the published article.

## A Appendix

### Formulars of the 6DOF-SVC Model

The 6DOF-SVC model [24], [34], [46], [57], see also [56], [58], uses transfer functions of the modeled semicircular channels (*SCC*) (eq. 5) and the otholite organ (*OTO*) (eq. 6) [30] as well as a Laplace function (*LP* ) (eq. 7) for sensory unification to capture the real movement. The semicircular canals detect angular velocity (*w*), and their inertial and filtering properties are modeled using a second-order transfer function. In contrast, the OTO senses linear acceleration (*f* ), including the gravitational component, which is mathematically incorporated into the physiological model without modification. In parallel, the perception and estimation of human’s own movement is estimated using an internal physiological reference model (*SCC, OTO* and *LP* (eq. 8, 9, 10), which is corrected by the difference to the movement actually perceived by the sense of balance (modeled by the inner ear: *OTO, SCC*).

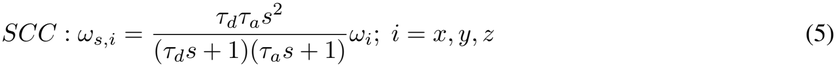

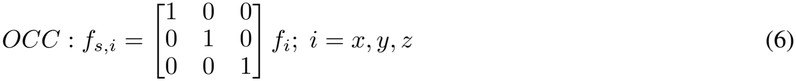

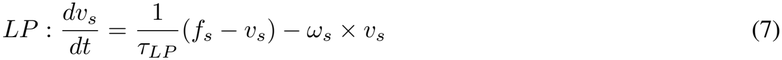

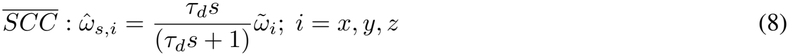

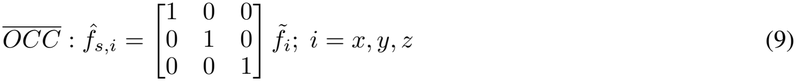

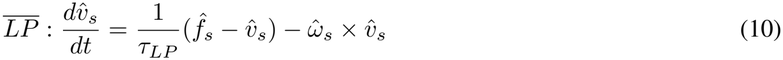

The estimated SC (Δ*v*) is then extracted and converted by a Hill-function to a normed one-dimensional signal (Δ*v*_*norm*_) (eq. 11). Subsequently, a PT2 element (eq. 12) simulates the MSI. Depending on the selected parameters and the intended application (e.g., as a target variable for model training), the MSI may represent different outcome measures, such as the probability of nausea [29], the probability of vomiting [32], [33], or a direct individual subjective rating (e.g., MISC scores, whereas the PT2 element did not represent the most suitable processing block for this approach) [37], [46].

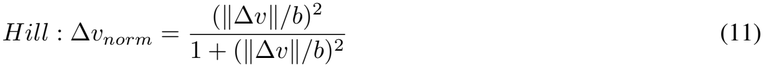

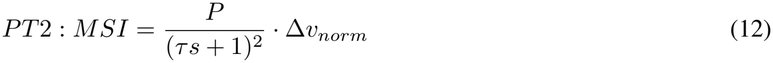

The parameters of the physiological functions (*τ*_*d*_ and *τ*_*a*_) are well established and will not be changed. Parameters *K*_*a*_ and *K*_*w*_ weight the input, while parameters *K*_*ac*_, *K*_*wc*_ and *K*_*vc*_ weight the loop gain, i.e. the SC. Hereby, *K*-parameters weight all three spatial dimensions (*x, y, z*) equally. The parameter *b* affects the contribution of the SC within the Hill function and, consequently, influences the gradient of the MSI. The parameter *τ* alters the temporal stability of the MSI, and the parameter *P* is the target value and maximum achievable MSI value of the PT2 element.

### Additional Information to the Experimental Driving Studies

S1: Vehicel & Route: Ford Galaxy 2.0 TDCi (WA6, 3^*rd*^ gen.) on a closed testing area (Fig. 9 (a)). Rear left seating position. NDRT: Reading and playing games (Simon Says, Connect Four, and Tic Tac Toe) in repetitive 5 minute sequences. Equipment: External IMU [VBOX IMU04, Racelogic Ltd, U.K.] placed behind co-driver seat, to collect acceleration and angular velocity with 100 Hz.

**Figure 9:**
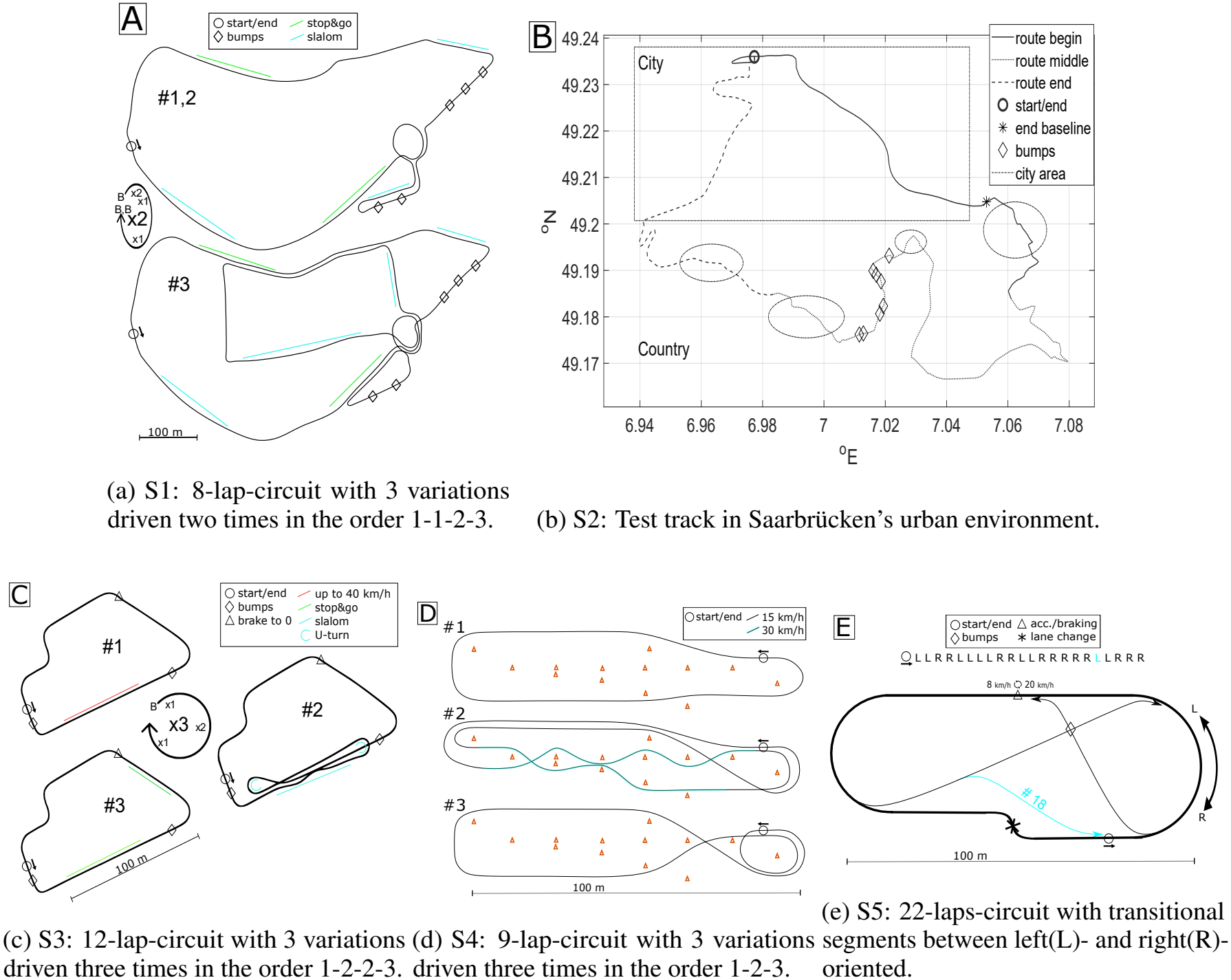
Test tracks of the five driving experiments.

**Figure 10:**
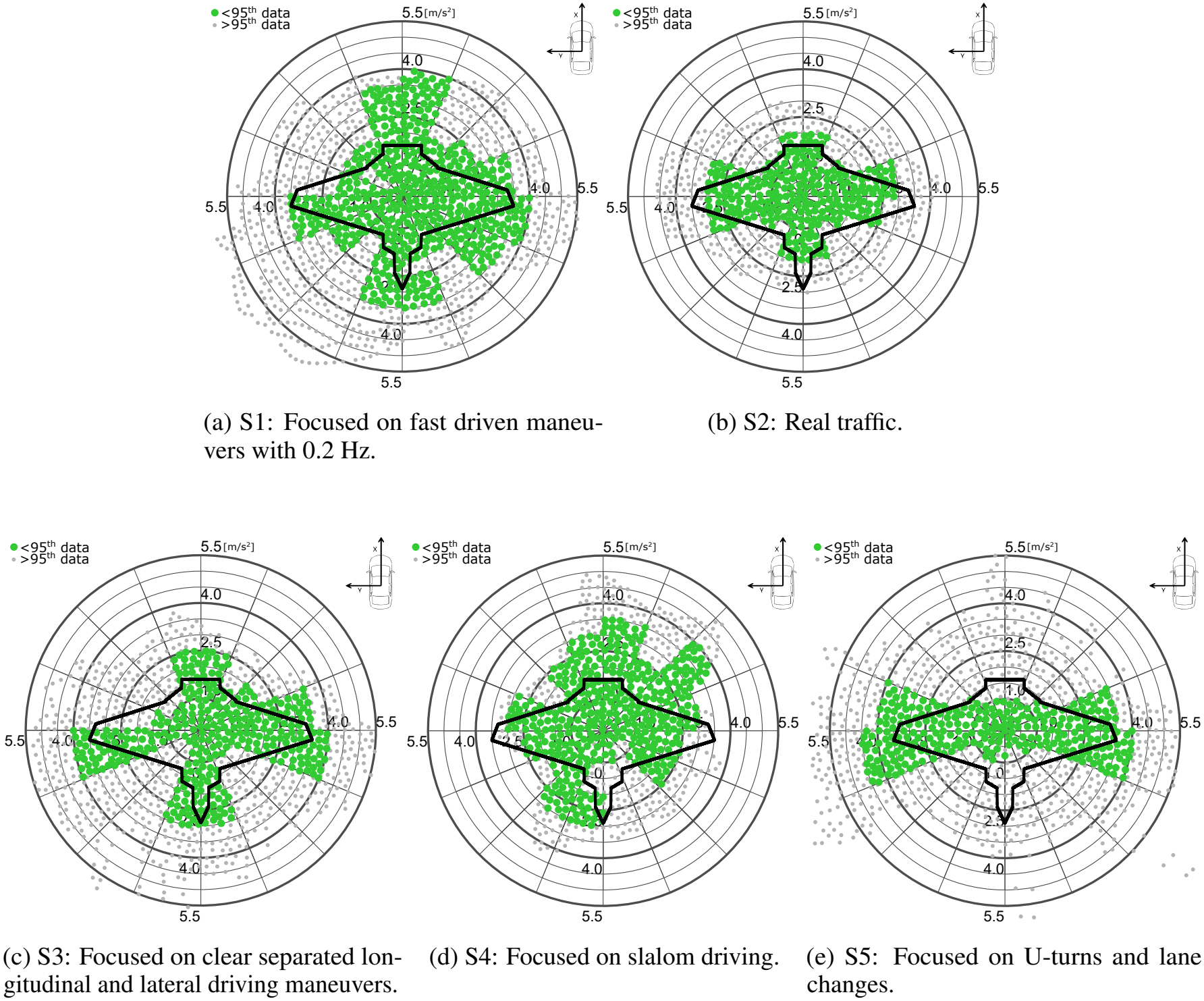
G–G-diagrams of the five experiments, with the 95th percentile data highlighted in green and the highest 5% in grey. A common reference driving profile is shown as a black line.

S2: Vehicel & Route: VW Touran (2^*nd*^ gen.) in real traffic (Fig. 9 (b)). Rear left seating position. NDRT: Reading, watching silent movie sequences and playing games (Simon Says and Tic Tac Toe) in repetitive 5 minute sequences. Equipment: Inbuilt sensors of the car to collect acceleration, angular Velocity and GPS with 100 Hz via CAN. External sensor unit [VBOX 3i + IMU04, Racelogic Ltd, U.K.] placed centrally beneath the center armrest, including with GPS/GLONASS dual antenna system to collect acceleration, angular velocity and GPS with 100 Hz.

S3: Vehicel & Route: VW Touran (2^*nd*^ gen.) on a closed testing area (Fig. 9 (c)). Rear center seating position. Countermeasure Systems: Within Subject Design (tactile system vs. (light) nonfunctional system) NDRT: Reading and watching documentary in repetitive 5 minute sequences. Equipment: Inbuilt sensors of the car to collect acceleration, angular Velocity and GPS with 80 Hz via CAN. External sensor unit [VBOX 3i + IMU04, Racelogic Ltd, U.K.] placed centrally beneath the center armrest, including GPS/GLONASS dual antenna system to collect acceleration, angular velocity and GPS with 80 Hz.

S4: Vehicel & Route: Two Audi Q5 2.0 TFSI (2^*nd*^ gen.) (one common car one with an active tilting suspension system) on a closed testing area (Fig. 9 (d)). Rear left seating position. Countermeasure Systems: Within Subject Design (active tilting vs. non active tilting) NDRT: Reading. Equipment: External INS [WTGAHRS2, WitMotion ShenZhen Co., Ltd.] placed centrally in the rear footwell, to collect acceleration, angular velocity and angles with 80 Hz.

S5: Vehicel & Route: VW Touran (2^*nd*^ gen.) on a closed testing area (Fig. 9 (e)). Rear center seating position. Countermeasure Systems: Between Subject Design (Sound system vs. light system vs. control group) NDRT: Reading. Equipment: Inbuilt sensors of the car to collect acceleration, angular Velocity and GPS with 80 Hz via CAN. External sensor unit [VBOX 3i + IMU04, Racelogic Ltd, U.K.] placed centrally beneath the center armrest, including GPS/GLONASS dual antenna system to collect acceleration, angular velocity and GPS with 80 Hz.

